# Dynamics of maternal gene expression in *Rhodnius prolixus*

**DOI:** 10.1101/2021.08.13.456198

**Authors:** Agustina Pascual, Rolando Rivera Pomar

## Abstract

The study of the developmental processes in *Rhodnius prolixus* has recently advanced with the sequencing of the genome. In this work, we study maternal gene expression driving oogenesis and early embryogenesis in *R. prolixus*. We analyze the transcriptional profile of mRNAs to establish the genes expressed across the ovary, unfertilized eggs and different embryonic stages of *R. prolixus* until the formation of the germ band anlage (0, 12, 24, and 48 hours post egg laying). We identified 81 putative maternal and ovary-related genes and validated their expression by qRT-PCR. Consistent with a role in oogenesis and early development of *R. prolixus*, we show that parental RNAi against *Rp-BicD* results in embryos that did not show any distinguishable embryonic structure. In this framework we propose three hierarchies of maternal genes that affect early and late oogenesis, and embryonic patterning.

## Introduction

During insect embryogenesis, a sequential series of dynamic processes that include cell division, growth and fate specification take place to establish the necessary components to give rise to a complete organism, playing a fundamental role to support the developmental process of the whole life cycle ^1-3^.

There are three modes of insect embryogenesis: long, intermediate, and short germ embryogenesis ^4^. Long germ embryogenesis is defined by the simultaneous establishment of all segmental fates at the blastoderm stage. This is a derived mode of embryogenesis, found in scattered species among the Holometabola, such as *Drosophila melanogaster*. These insects have polytrophic meroistic ovaries. In short or intermediate germ insects only the most anterior segments are specified before gastrulation, while the more posterior segments are generated and patterned progressively from a posterior region called the growth zone. This represents an ancestral type of insect embryogenesis, described in insect models such *Oncopeltus fasciatus, Rhodnius prolixus, Bombyx mori, Tribolium castaneum*, It corresponds to insects with telotrophic or panoistic ovaries ^4-8^. A common feature across the different modes of embryogenesis is the loading of maternal mRNA transcripts and proteins in the egg during oogenesis ^9^.

In the last 20 years the rise of new models for comparative insect development provided a framework to understand the genetic basis of development and evolution ^10-17^. The different mechanisms of insect embryogenesis are determined by specific spatiotemporal gene expression patterns derived from common genetic program, suggesting that the mechanisms are much more conserved than the diversity of germ types might suggest ^4,18-22^. In addition, a detailed study of cell flow during germ band extension and the fate map of *T. castaneum* embryo led to the idea that short and long germ bands share many more common features than thought ^23^. In the last decade the expanse of genomics and transcriptomic analysis provided an insight of the transcriptional basis of the embryonic development in non-model insect species ^24^. However, the complete repertory of genes involved in oogenesis and early embryogenesis has been reported in detail only in *D. melanogaste*r ^25-29^ and *T. castaneum* ^15,30,31^, remaining an open question in other model organisms.

The blood-feeding insect *Rhodnius prolixus* is one vectors of *Trypanosoma cruzi*, the etiologic agent of Chagas disease ^32,33^. In addition to its medical interest, it has been a classical model for physiology and biochemistry ^34-37^. The embryonic development of *R. prolixus* has been described from fertilization to hatching ^7^, and the process of oogenesis studied in detail ^38-44^. The genome was recently sequenced ^45^ and since then, *R. prolixus* is an emerging model for developmental biology ^46-50^. Several transcriptome analyses were reported, focusing in the gene expression of the follicular epithelium, the early previtellogenic stage of oogenesis; as well, in the impact of the nutritional state on regulatory pathways associated with reproductive performance ^51-53^. Very recently, a thorough study on previtellogenic ovaries and unfertilized eggs discovered a large number of unannotated genes in the *R. prolixus* genome and unveiled a large set of maternal genes ^54^. With all this knowledge in place, we have moved forward to the understanding of the genetic and molecular mechanisms driving oogenesis and the maternal contribution to embryo patterning in *R. prolixus*. Here, we present a transcriptome profiling approach to identify the genetic basis underlying oogenesis and early embryogenesis of *R. prolixus* until the onset of gastrulation, with a focus on genes related to embryonic patterning and egg formation. We provide novel insight into the molecular basis of early embryo formation and show the dynamic of mRNA expression during early embryo development in *R. prolixus*. Our study provides maternal and early embryonic transcriptomes of this hemimetabolous insect. We present a comprehensive qualitative data about genes related to segmentation, dorsal ventral axis and oogenesis, validate gene expression by qRT-PCR and show the phenotype of *Bicaudal D* (*BicD*) homolog likely related to early steps of the maternal cascade that leads to patterning.

## Materials and methods

### Insect rearing

A colony of *R. prolixus* was maintained in our laboratory in a 12:12 hour light/dark period at 28 °C and 80% relative humidity in controlled environment incubators. In these conditions, embryogenesis takes 14 ± 1 days. Insects were regularly fed on chickens, which were housed, cared, fed and handled in accordance with resolution 1047/2005 (Consejo Nacional de Investigaciones Científicas y Técnicas, CONICET) regarding the bioethical framework for biomedical research with laboratory, farm, and nature collected/wild animals. This framework is in accordance with international standard procedures. Biosecurity considerations agree with CONICET resolution 1619/2008, which is in accordance with the WHO Biosecurity Handbook (ISBN 92 4 354 6503).

### Sample collection, RNA isolation and sequencing

Adult mated insects 6^th^ days after the feeding regimen were used to collect fertilized eggs at specific points in developmental time – 0 (zygote), 12 (blastoderm), 24 (cellular blastoderm) and 48 (onset of germ band formation) hours post egg laying (*hPL*). Virgin female adults were used to collect unfertilized eggs, which were immediately frozen and stored in liquid nitrogen. At the same time, female ovaries were dissected in the vitellogenic stage and placed in a cryotube containing Trizol (Invitrogen), flash frozen and stored in liquid nitrogen until use.

For the transcriptome profiling, RNA was extracted from 150 embryos for each developmental time and 30 vitellogenic ovaries. For qRT-PCR analysis, independent experiments were carried out using 75 embryos from each specific time and 10 vitellogenic ovaries. Total RNA was isolated using Trizol (Invitrogen) as recommended by the manufacturer. RNA integrity was determined by agarose electrophoresis and concentration measured using Qubit RNA Assay Kit in a Qubit 2.0 Fluorometer (Life Technologies, Invitrogen). cDNA libraries were synthetized from 1 µg of total RNA and sequenced using a HiSeq-3000 platform (Illumina) to obtain the 50 base pairs (bp) (single-end) or 150 bp (paired-end) reads. The RNA-seq data has been submitted to the NCBI SRA database, available under accession code PRJNA694974.

### Quality control, alignment and transcriptome assembly

Raw data were processed with FASTX-toolkit software (http://hannonlab.cshl.edu/fastx_toolkit/), to remove adapter sequences, reads with unknown bases and reads with quality scores lower than Q30, showed by the FastQC report (http://www.bioinformatics.babraham.ac.uk/projects/fastqc/). To avoid contaminants, the removal of adaptor sequences was ruled out using BLASTn ^55^ and the UniVec database (ftp://ftp.ncbi.nlm.nih.gov/pub/UniVec/) from NCBI. Additionally, remove rRNA sequences the SILVA database was used ^56^. The remaining reads were defined as clean reads and used for subsequent bioinformatics analyses. Tophat2 ^57^ was used to map clean reads to the *ab initio* annotations of *R. prolixus*, genome dataset version RproC3.3 ^45,58^. The mapping statistics by the RNA-seq reads were calculated by using bam_stat.py implemented in the RSeQC package ^59^ and the advanced statistics of coverage analysis were performed by the Qualimap application ^60^. After Tophat alignment, transcripts were assembled using Cufflinks ^61,62^. Assembly quality was assessed for each assembly using BUSCO analysis ^63^, with the reference gene set of arthropod (2.676 proteins) with default parameters. Fasta Statistics was used to display summary statistics from each transcriptome generated ^64^. The eggNOG 5.0 database was used for functional annotation of the transcripts with common denominators or functional categories (i.e., derived from the original COG categories). Also, predicted protein-coding transcripts ^65^ were functionally annotated. For each protein sequence protein signatures were assigned, using InterProScan search Version 5.0.0, ^66^ through the PfamA and SuperFamily databases. Proteins annotated by signatures were assigned into GO (Gene Ontology) categories, including biological processes (BP), molecular functions (MF) and cellular components (CC). To statistically analyze GO-term enrichment, topGO package ^67^ was implemented, using Fisher’s exact test and the false discovery rate (FDR) adjusted method. A q-value smaller than 0,05 were considered as significant. The reference set of gene-to-GO mappings was available from VectorBase (https://www.vectorbase.org/).

### Oogenesis and early embryogenesis gene identification

Gene identification was performed using local BLAST ^55^ on the six transcriptome assemblies. The BLAST algorithm used was BLASTx. The search was limited to 84 protein sequences derived from FlyBase (Version FB2020_03, https://flybase.org/), comprising genes related to oogenesis and early embryogenesis, with an e-value threshold of 10^−5^. Transcript with blast hit to *Drosophila* were then manually checked by BLASTx against all Arthropoda protein sequences (NCBI non-redundant protein (nr) database, assessed January 2018) to establish orthology. BLAST results were classified into the known *D. melanogaster* developmental process. For the maternal gene search, a database was generated from different resources containing 10.277 specific protein sequences (Additional file 1) ^15,25-29^.

### Quantitative real-time PCR

Total RNA was isolated using Trizol reagent (Invitrogen) and treated with DNAse (QIAGEN). cDNA was synthesized using SuperScript™ VILO™ MasterMix kit (Invitrogen) following the manufacturer’s instructions. PCR was performed in technical triplicates (3 wells/cDNA sample), in a 10 μl final volume as follows: (i) 95 °C for 10 min; (ii) 95 °C for 15 sec; (iii) 55 °C for 30 sec; (iv) 72 °C for 45 sec; (v) steps (ii) to (iv) for 35 cycles. Gene expression level was quantified using SsoAdvanced Universal SYBR Green Supermix (Bio-Rad) in an Applied Biosystems 7.500 Real-Time PCR System (Thermo Fisher Scientific). A control without a template was included in all batches and *α-tubulin* was used as reference, after a screen of several housekeeping gene candidates, as it provided consistent results on the embryonic stages analyzed. All primer pairs (Additional file 2) were tested for dimerization, efficiency, and amplification of a single product. The Ct value was averaged for the technical triplicate experiments and subtracted from the average Ct of the reference gene, to yield the expression difference (dCt) for each biological replicate. The results were analyzed according to ^68^. To test whether the expression of a given gene was significantly different across developmental times, a one-way ANOVA was carried out followed by post-hoc test using GraphPad Prism v6.0 software (GraphPad Software, CA, USA, www.graphpad.com).

### *In situ* Hybridization and Parental RNAi

DNA templates used to synthesize *in situ* hybridization probes were obtained by PCR using oligonucleotides carrying T7 promoter sequences at the 5’-end. The templates were *in vitro* transcribed using DIG RNA labeling kit (Roche). *In situ* hybridization was performed as described in Pascual, et al. ^50^.

For parental RNAi, dsRNA was produced by simultaneous transcription with T7 RNA polymerase (New England Biolabs) on templates containing T7 promoter sequences at both ends. dsRNA^*BicD*^ was quantitated and injected into virgin females as described in Lavore, et al. ^69^. Two days after injection, the females were fed to induce oogenesis and mated. After mating, eggs were collected and ovaries fixed as described ^50^. A negative control was performed injecting virgin females with dsRNA corresponding to the β- lactamase gene (dsRNA^β- lac^) of *E. coli* ^69^. Amplicon was sequenced to confirm identity (Macrogen Inc.). Oligonucleotides used in this study are listed in the Additional file 2.

## Results and discussion

### 1. Assembling the ovarian and early embryonic transcriptomes of *R. prolixus*: characterization and completeness analysis

The RNA-seq output comprised six *R. prolixus* samples that cover late oogenesis to the beginning of germ band extension (48 *hPL*). Statistics on the sequencing and mapping are reported in Table 1A. According to completeness analysis, the coverage metrics obtained indicated that the assembled transcriptomes are sufficient for a meaningful analysis (Table 1B). As the genomic reference has a reasonable amount of positions that are not called, transcriptomes assembled by mapping genomic predictions (*ab initio*) were used to subsequent analyzes. In this respect, a review of zygotic genes ^14^ and of the regulatory pathways involved in egg production ^52^ has been reported based on these genomic annotations with a robust gene identification.

**Table 1.**
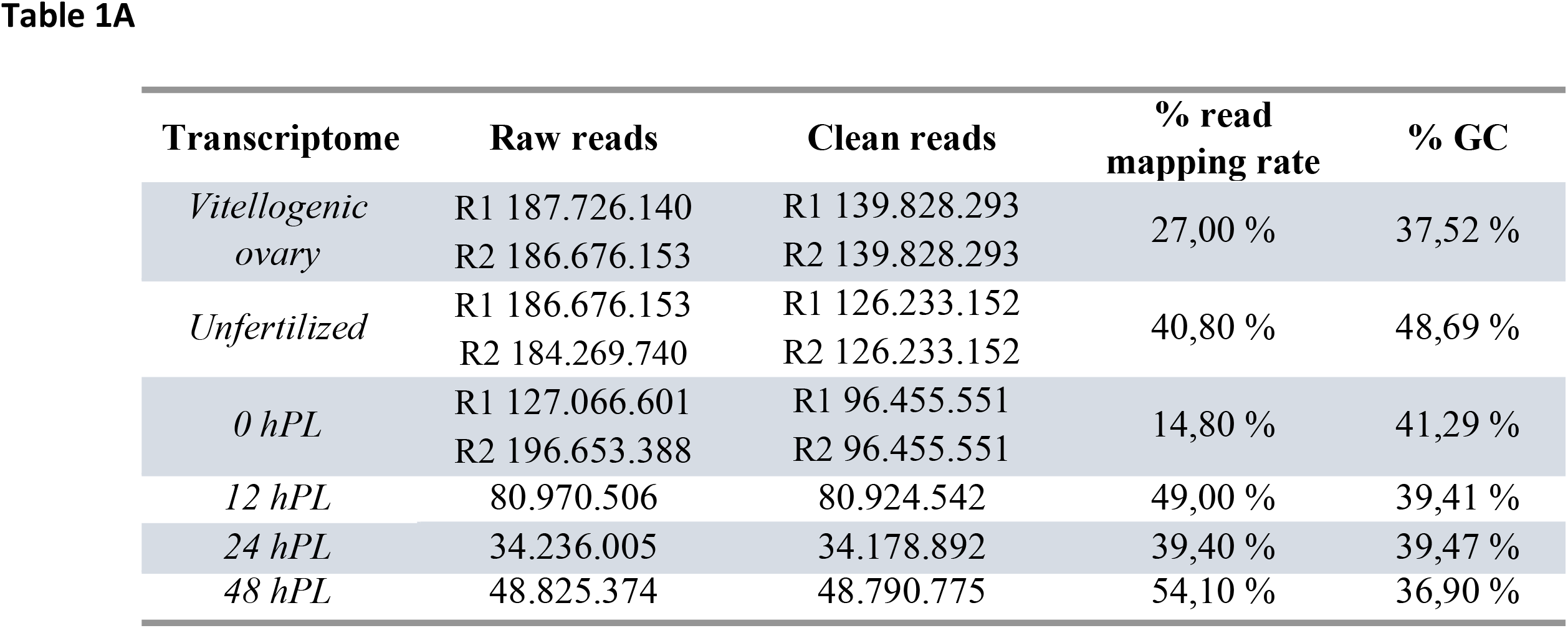
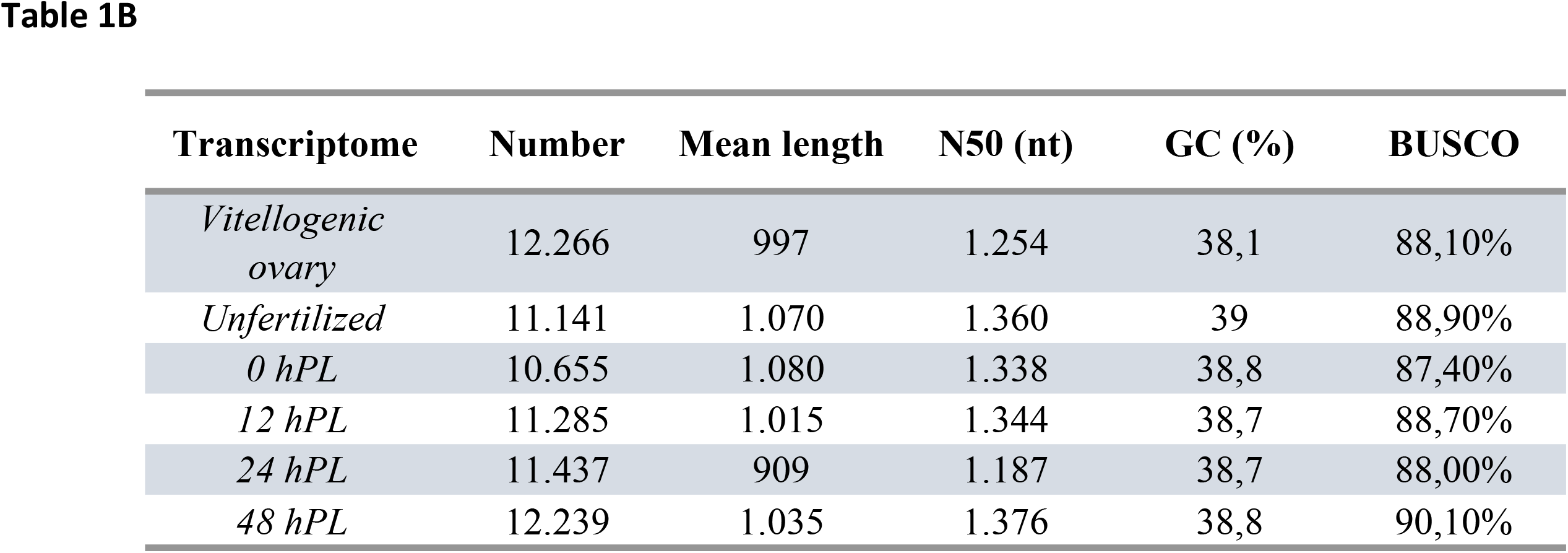
Summary of the RNA-seq metrics from *R. prolixus* transcriptomes from vitellogenic ovaries, unfertilized and 0 to 48 *hPL* eggs. **A**: Raw Reads: the original sequencing reads counts; Clean Reads: number of reads after filtering; % read mapping rate: percentages of the overall mapping rate using the annotated genome as reference, % GC: percentages of G and C in total bases. **B**: Transcriptome assembly’s statistics, Number, number of the total of reconstructed transcripts; Mean length: the mean length in base pairs; N50: is the size of the transcript which, along with the larger transcripts, contain half of sequence of the reference; GC: percentages of G and C in total bases; BUSCO: percentages of the transcriptome completeness by BUSCO analysis results.

In order to conduct a transcriptome-composition representation analysis, eggNOG analysis was performed. A total of 25 eggNOG categories were detected (Figure 1 and Additional file 3), in which the category “function unknown” was dominant followed by “Signal transduction mechanisms” and “Post-translational modification, protein turnover, and chaperones” in all the analyzed developmental times. To obtain information of the predicted proteins, InterproScan searched were performed to identified functional domains, repeats, sites and protein families conserved in the protein-coding transcripts (Table 2 and Additional file 4). For all of the characterized transcripts, statistically over-represented GO terms were identified using the FDR adjusted relative to a reference set of 11.947 genes. These statistically highlighted GO terms were summarized to generic GO categories for each developmental time studied (Table 2 and Additional file 5). GO analysis showed that mainly metabolic processes were enriched during embryo development, such as cellular macromolecule metabolic process (GO: 0044260), nucleobase-containing compound metabolic process (GO:0006139), organonitrogen compound biosynthetic process (GO:1901566), gene expression (GO: 0010467). This enrichment is in agreement with the requirements of the embryo during the transitions between the different embryonic stages with rapidly changing of the anabolic and catabolic demands ^70,71^.

**Figure 1.**
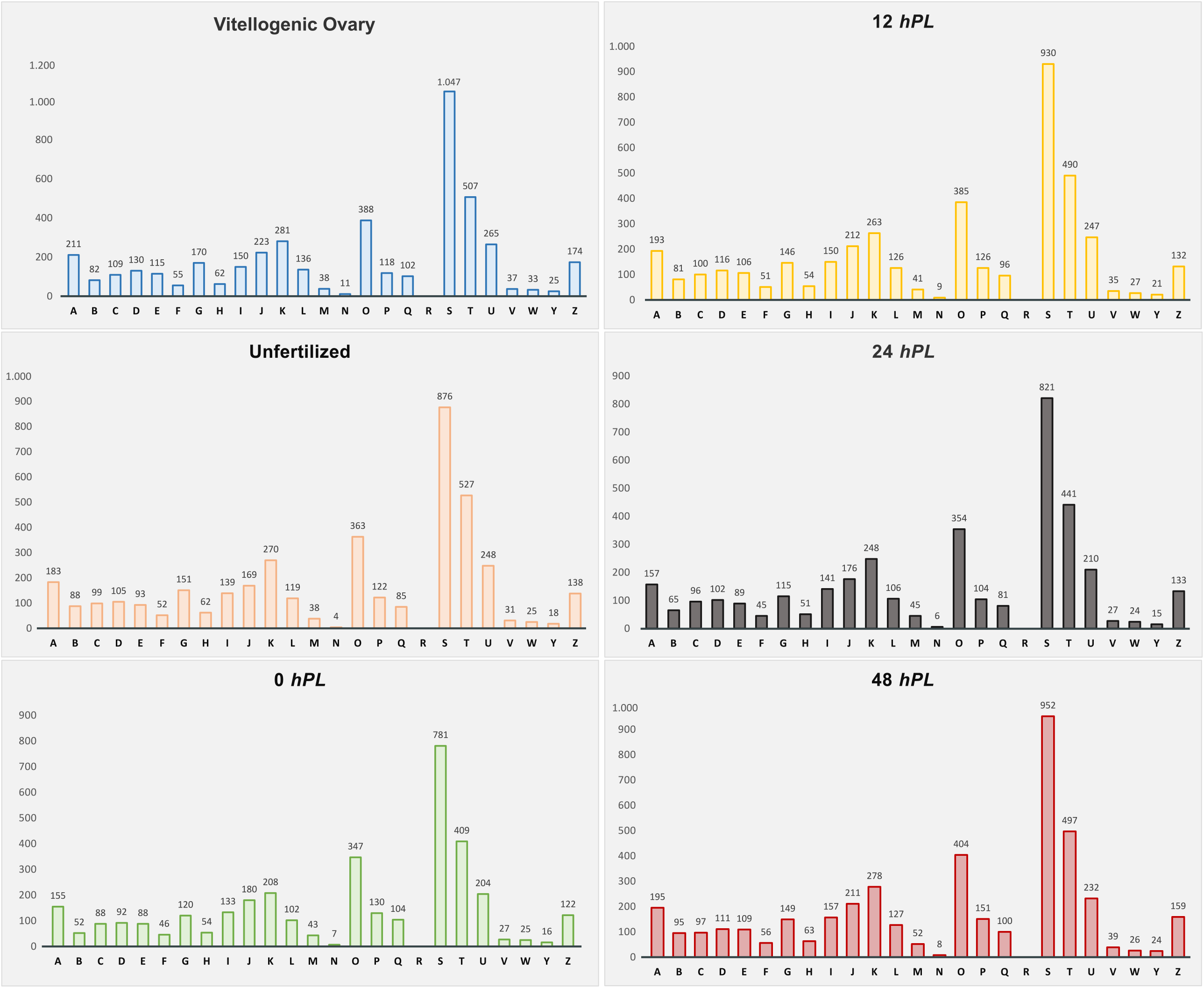
Classification of eggNOG annotations in the *R. prolixus* transcriptomes. The capital letters on the X-axis represent different eggNOG categories. Y-axis shows the number of transcripts in each eggNOG category. A: “RNA processing and modification”, B: “Chromatin structure and dynamics”, C: “Energy production and conversion”, D: “Cell cycle control, cell division, chromosome partitioning”, E: “Amino acid transport and metabolism”, F: “Nucleotide transport and metabolism”, G: “Carbohydrate transport and metabolism”, H: “Coenzyme transport and metabolism”, I: “Lipid transport and metabolism”, J: “Translation, ribosomal structure and biogenesis”, K: “Transcription”, L: “Replication, recombination and repair”, O: “Post-translational modification, protein turnover, and chaperones”, P: “Inorganic ion transport and metabolism”, Q: “Secondary metabolites biosynthesis, transport, and catabolism”, R: “General function prediction only”, S: “Function unknown”, T: “Signal transduction mechanisms”, U: “Intracellular trafficking, secretion, and vesicular transport”, V: “Defense mechanisms”, W: “Extracellular structures”, Y: “Nuclear structure”, Z: “Cytoskeleton”.

**Table 2.** Summary of the functional annotation. Top 10 Interpro domains and GO terms enriched in each transcriptome analyzed. TopGO results were shown in different colors according to the three sub-ontologies: green “Biological Process”, blue “Molecular functions” and orange “Cellular component”.

| Transcriptome | InterPro analysis |  |  | TopGO enrichment analysis |  |  |
| --- | --- | --- | --- | --- | --- | --- |
|  | Number accession | Description | Number of transcripts | GO ID | GO Term | q-value |
| *Vitellogenic Ovary* | IPR001680 | WD40 repeat | 143 | GO:0044260 | cellular macromolecule metabolic process | 2,00E-23 |
|  | IPR027417 | P-loop containing nucleoside triphosphate hydrolase | 71 | GO:0034641 | cellular nitrogen compound metabolic process | 3,60E-23 |
|  | IPR016024 | Armadillo-type fold | 49 | GO:0044237 | cellular metabolic process | 1,60E-21 |
|  | IPR036236 | Zinc finger C2H2 superfamily | 48 | GO:0010467 | gene expression | 2,45E-18 |
|  | IPR000504 | RNA recognition motif domain | 46 | GO:0003723 | RNA binding | 2,20E-09 |
|  | IPR035979 | Zinc finger C2H2-type | 44 | GO:0003735 | structural constituent of ribosome | 1,80E-06 |
|  | IPR013087 | RNA-binding domain superfamily | 44 | GO:0016740 | transferase activity | 2,40E-04 |
|  | IPR036322 | WD40-repeat-containing domain superfamily | 42 | GO:0005622 | intracellular | 4,00E-24 |
|  | IPR036249 | Thioredoxin-like superfamily | 42 | GO:0044424 | intracellular part | 4,00E-24 |
|  | IPR011990 | Tetratricopeptide-like helical domain superfamily | 38 | GO:0005623 | cell | 4,00E-24 |
| *Unfertilized* | IPR001680 | WD40 repeat | 140 | GO:0044260 | cellular macromolecule metabolic process | 3,60E-22 |
|  | IPR027417 | P-loop containing nucleoside triphosphate hydrolase | 69 | GO:0034641 | cellular nitrogen compound metabolic process | 4,00E-21 |
|  | IPR036236 | Zinc finger C2H2 superfamily | 51 | GO:0044237 | cellular metabolic process | 1,07E-18 |
|  | IPR016024 | Armadillo-type fold | 47 | GO:0010467 | gene expression | 4,40E-18 |
|  | IPR011990 | Tetratricopeptide-like helical domain superfamily | 46 | GO:0003723 | RNA binding | 2,40E-08 |
|  | IPR000504 | RNA recognition motif domain | 44 | GO:0003735 | structural constituent of ribosome | 1,10E-06 |
|  | IPR036249 | Thioredoxin-like superfamily | 43 | GO:0005622 | intracellular | 4,00E-24 |
|  | IPR013087 | Zinc finger C2H2-type | 42 | GO:0044424 | intracellular part | 4,00E-24 |
|  | IPR036322 | WD40-repeat-containing domain superfamily | 40 | GO:0032991 | protein-containing complex | 4,00E-24 |
|  | IPR035979 | RNA-binding domain superfamily | 39 | GO:0005623 | cell | 4,00E-24 |
| *0 hPL* | IPR001680 | WD40 repeat | 134 | GO:0044260 | cellular macromolecule metabolic process | 3,60E-22 |
|  | IPR027417 | P-loop containing nucleoside triphosphate hydrolase | 69 | GO:0034641 | cellular nitrogen compound metabolic process | 4,00E-21 |
|  | IPR036236 | Zinc finger C2H2 superfamily | 55 | GO:0044237 | cellular metabolic process | 1,07E-18 |
|  | IPR013087 | Zinc finger C2H2-type | 49 | GO:0010467 | gene expression | 4,40E-18 |
|  | IPR011990 | Tetratricopeptide-like helical domain superfamily | 48 | GO:0003723 | RNA binding | 2,40E-08 |
|  | IPR016024 | Armadillo-type fold | 47 | GO:0003735 | structural constituent of ribosome | 1,10E-06 |
|  | IPR000504 | RNA recognition motif domain | 42 | GO:0005622 | intracellular | 4,00E-24 |
|  | IPR036249 | Thioredoxin-like superfamily | 41 | GO:0044424 | intracellular part | 4,00E-24 |
|  | IPR036322 | WD40-repeat-containing domain superfamily | 40 | GO:0032991 | protein-containing complex | 4,00E-24 |
|  | IPR015919 | Cadherin-like superfamily | 40 | GO:0005623 | cell | 4,00E-24 |
| *12 hPL* | IPR001680 | WD40 repeat | 142 | GO:0044260 | cellular macromolecule metabolic process | 2,00E-23 |
|  | IPR027417 | P-loop containing nucleoside triphosphate hydrolase | 71 | GO:0034641 | cellular nitrogen compound metabolic process | 4,00E-23 |
|  | IPR036236 | Zinc finger C2H2 superfamily | 54 | GO:0044237 | cellular metabolic process | 5,20E-22 |
|  | IPR013087 | Zinc finger C2H2-type | 48 | GO:0010467 | gene expression | 6,50E-19 |
|  | IPR016024 | Armadillo-type fold | 47 | GO:0003723 | RNA binding | 1,18E-08 |
|  | IPR000504 | RNA recognition motif domain | 45 | GO:0003735 | structural constituent of ribosome | 1,40E-06 |
|  | IPR036322 | WD40-repeat-containing domain superfamily | 42 | GO:0016740 | transferase activity | 3,47E-05 |
|  | IPR035979 | RNA-binding domain superfamily | 42 | GO:0140098 | catalytic activity, acting on RNA | 1,85E-03 |
|  | IPR011990 | Tetratricopeptide-like helical domain superfamily | 36 | GO:0005622 | intracellular | 4,00E-24 |
|  | IPR011009 | Protein kinase-like domain superfamily | 35 | GO:0044424 | intracellular part | 4,00E-24 |
| *24 hPL* | IPR001680 | WD40 repeat | 137 | GO:0044260 | cellular macromolecule metabolic process | 2,00E-23 |
|  | IPR027417 | P-loop containing nucleoside triphosphate hydrolase | 71 | GO:0034641 | cellular nitrogen compound metabolic process | 1,90E-22 |
|  | IPR036236 | Zinc finger C2H2 superfamily | 48 | GO:0044237 | cellular metabolic process | 8,00E-22 |
|  | IPR000504 | RNA recognition motif domain | 47 | GO:0010467 | gene expression | 9,00E-18 |
|  | IPR016024 | Armadillo-type fold | 45 | GO:0003723 | RNA binding | 5,60E-08 |
|  | IPR035979 | RNA-binding domain superfamily | 42 | GO:0003735 | structural constituent of ribosome | 1,80E-06 |
|  | IPR013087 | Zinc finger C2H2-type | 41 | GO:0016740 | transferase activity | 1,73E-05 |
|  | IPR036322 | WD40-repeat-containing domain superfamily | 40 | GO:0005622 | intracellular | 4,00E-24 |
|  | IPR011990 | Tetratricopeptide-like helical domain superfamily | 38 | GO:0044424 | intracellular part | 4,00E-24 |
|  | IPR011009 | Protein kinase-like domain superfamily | 37 | GO:0044464 | cell part | 4,00E-24 |
| *48 hPL* | IPR001680 | WD40 repeat | 147 | GO:0044260 | cellular macromolecule metabolic process | 2,00E-23 |
|  | IPR027417 | P-loop containing nucleoside triphosphate hydrolase | 74 | GO:0034641 | cellular nitrogen compound metabolic process | 3,10E-23 |
|  | IPR016024 | Armadillo-type fold | 47 | GO:0044237 | cellular metabolic process | 8,00E-20 |
|  | IPR011990 | Tetratricopeptide-like helical domain superfamily | 46 | GO:0010467 | gene expression | 7,50E-19 |
|  | IPR036249 | Zinc finger C2H2 superfamily | 45 | GO:0003723 | RNA binding | 6,80E-09 |
|  | IPR036236 | Thioredoxin-like superfamily | 45 | GO:0003735 | structural constituent of ribosome | 4,30E-06 |
|  | IPR000504 | RNA recognition motif domain | 45 | GO:0016740 | transferase activity | 2,20E-04 |
|  | IPR036322 | WD40-repeat-containing domain superfamily | 42 | GO:0005622 | intracellular | 4,00E-24 |
|  | IPR035979 | RNA-binding domain superfamily | 42 | GO:0044424 | intracellular part | 4,00E-24 |
|  | IPR013087 | Zinc finger C2H2-type | 40 | GO:0005623 | cell | 4,00E-24 |

A total of 1.192 annotated transcripts common to all developmental stages studied were further examined to determine GO enrichment. The represented GO terms (Figure 2 and Additional file 6) were categorized in two main groups: cellular components and biological processes. The main terms of cellular components are cell structures involved in protein synthesis, and the biological processes significantly over-represented terms are related to lipid, carbohydrate, nucleic acid and protein metabolism. As expected in the enrichment observed for each developmental time, these biological processes play key roles in the embryonic development of *R. prolixus*. The energy is supplied to the embryogenesis to proceed through the breakdown of biomolecules stored in the yolk ^72^. These, in turn, drive converging biosynthetic pathways such as protein and nucleotide biosynthesis required to meet the needs of the developing embryo ^70^. These results agree with the increment of lipid, protein and carbohydrate reported during embryonic stages of *T. castaneum, Boophilus microplus* and *Aedes aegypti* ^73-75^; and in fed females in *R. prolixus* and also other triatomines ^76-79^.

**Figure 2.**
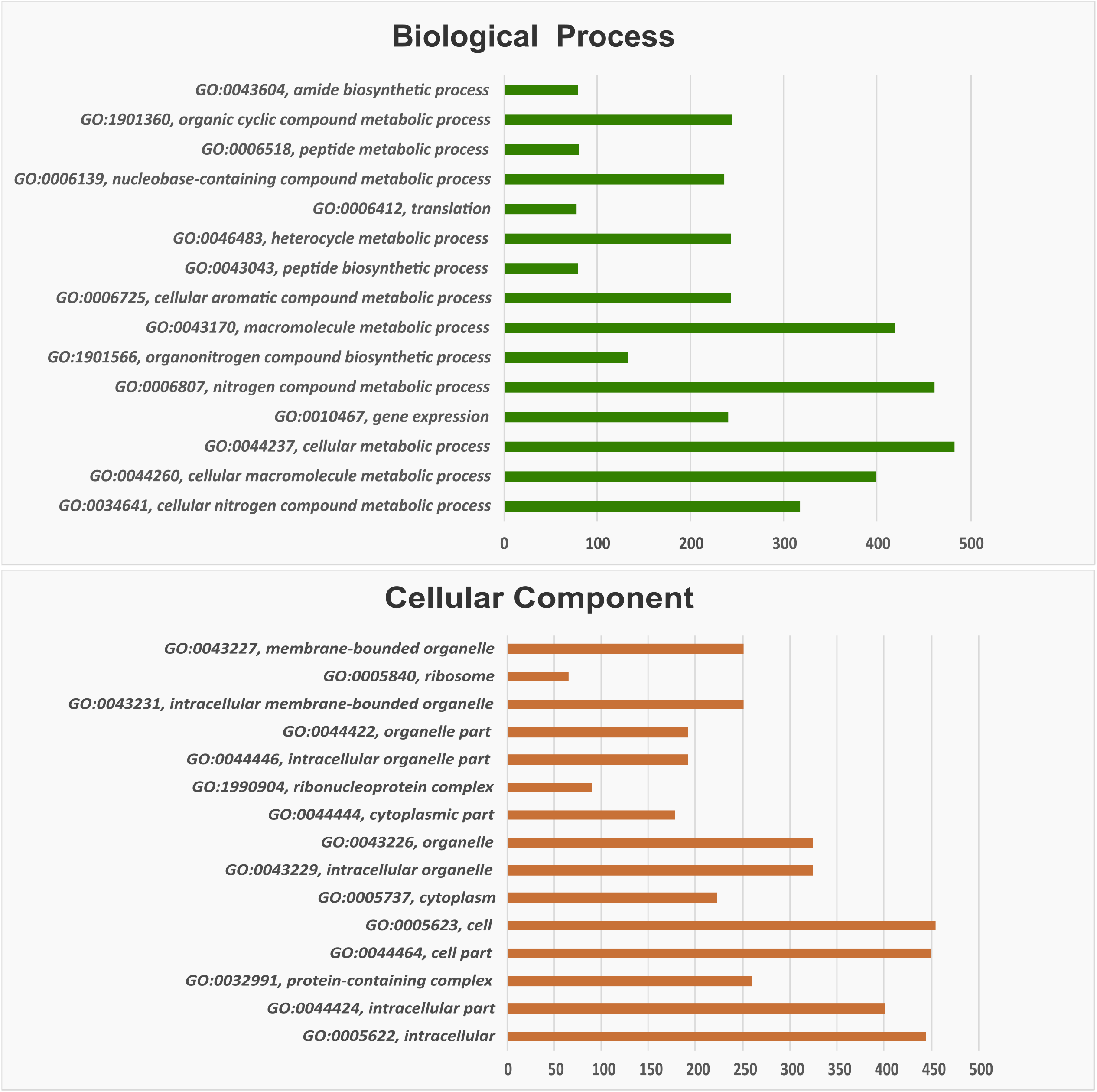
Bar graph of Gene Ontology (GO) enrichment analysis of the common transcripts across the six developmental stages. Upper: GO category “Biological process”, lower: “Cellular component”. X-axis: number of transcripts involved in the distinct GO terms. Y-axis: description of GO terms with the GO ID.

### 2. Gene identification for developmental processes

In order to study genes involved in early embryogenesis and oogenesis, a list of 84 developmental genes (37 segmentation-related genes, 16 of the dorsal patterning pathway and 31 linked to oogenesis) related to these processes in *D. melanogaster* was used to identified orthologues sequences in the *R. prolixus* transcriptomes. (Table 3A-C).

**Table 3.**
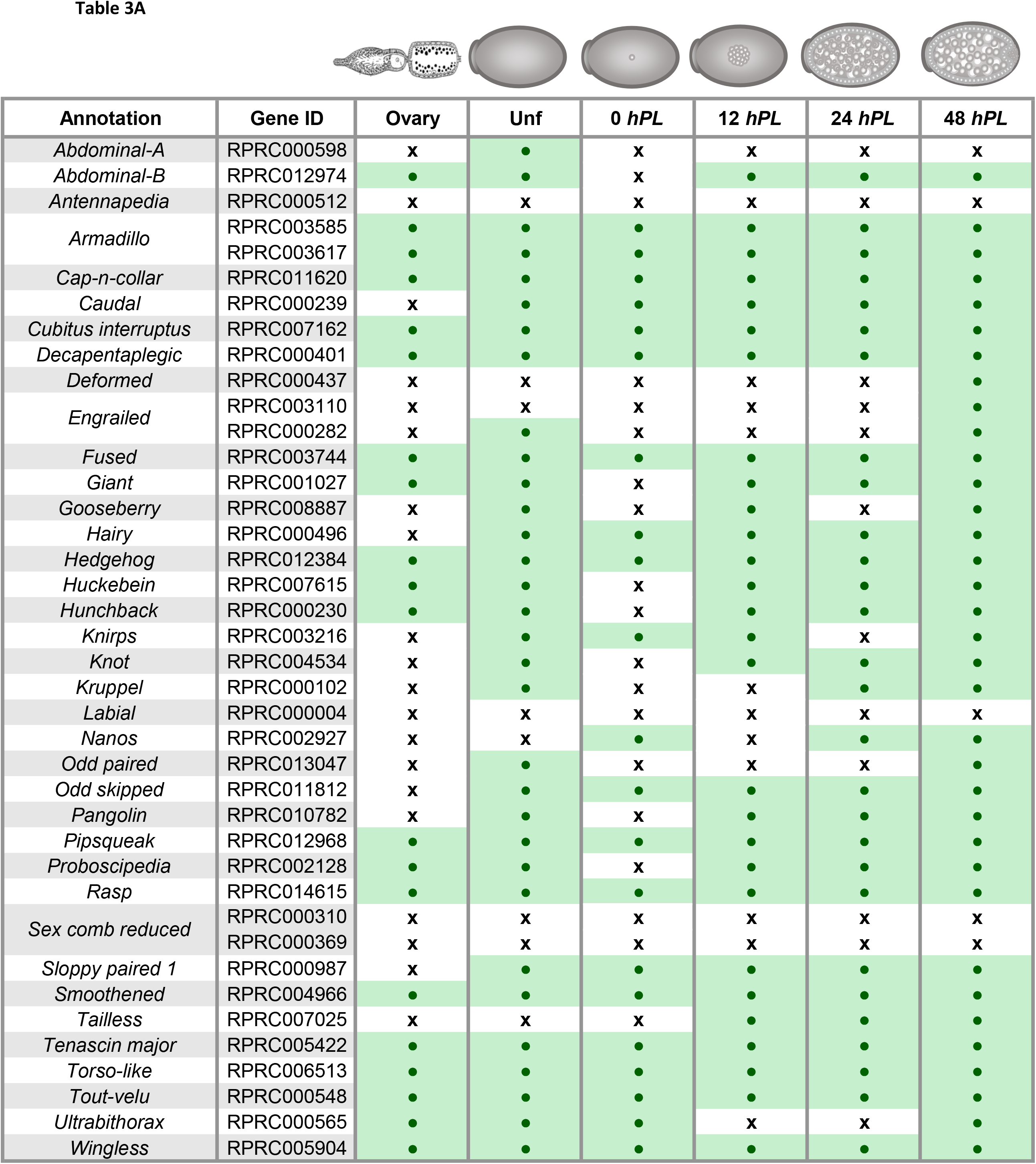
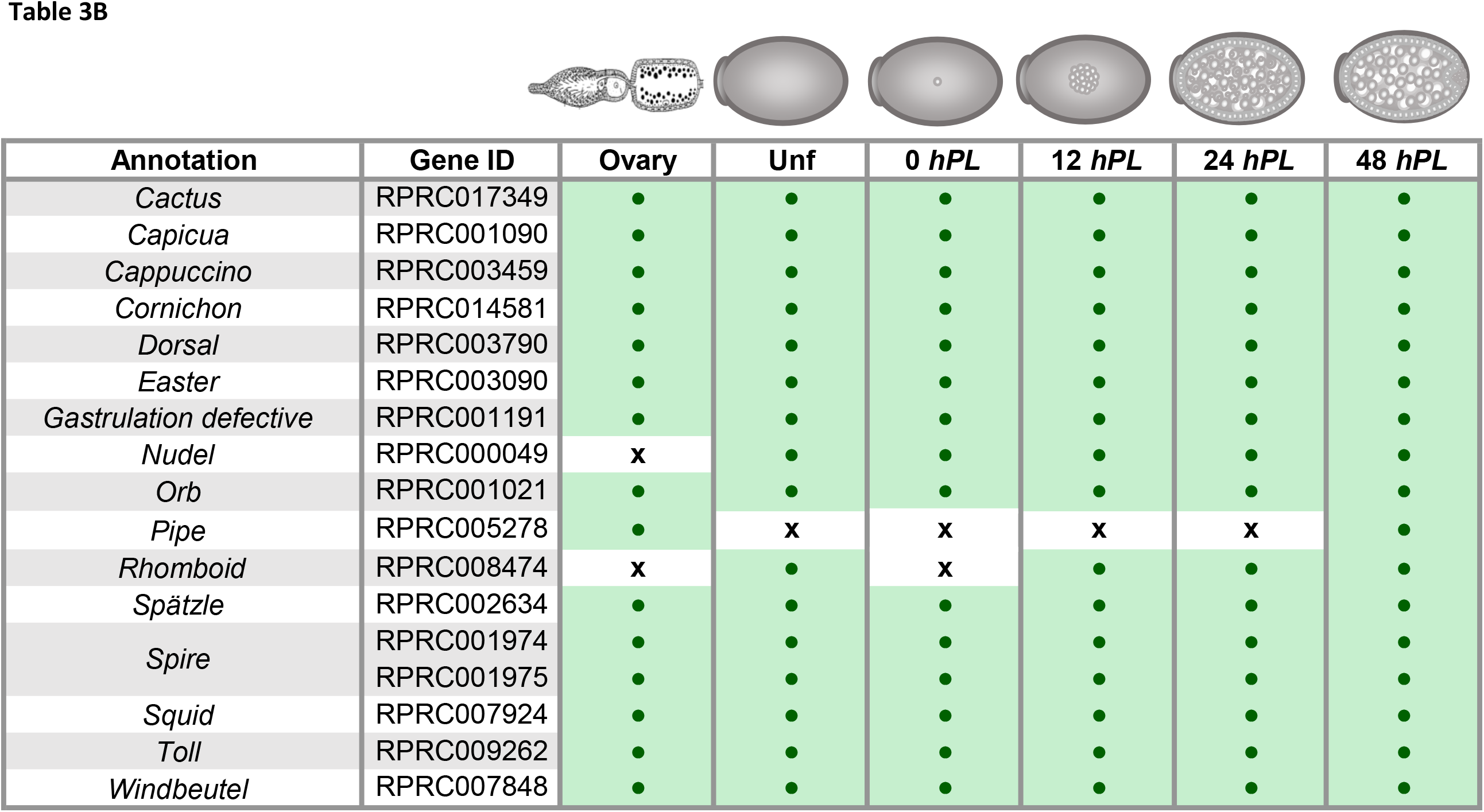
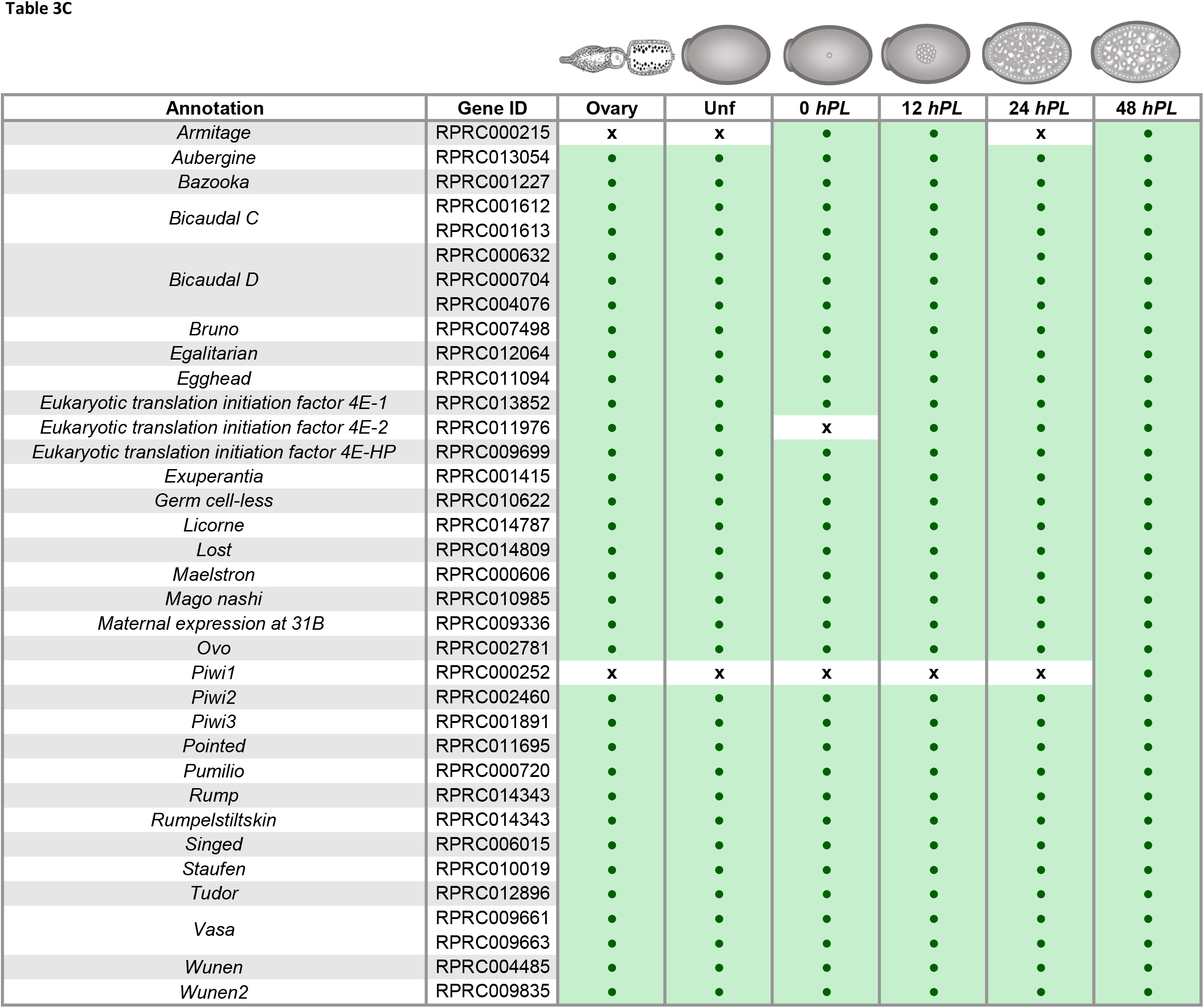
Developmental genes identified in the *R. prolixus* transcriptomes. Gene ID: VectorBase code (the official gene number in the RproC3 genome assembly); Annotations: protein name we are assigning. For each annotation, presence/absence was assessed by sequence similarity search. Green boxes indicate presence of expression; white boxes indicate absence of expression. **A:** Segmentation genes, **B:** Dorso-ventral genes, **C:** oogenesis, orthologues *piwi* genes were annotated as Brito, et al. ^48^.

| Annotation | Gene ID | Ovary | Unf | 0 hPL | 12 hPL | 24 hPL | 48 hPL |
| --- | --- | --- | --- | --- | --- | --- | --- |
| Abdominal-A | RPRC000598 | x | ● | x | x | x | x |
| Abdominal-B | RPRC012974 | ● | ● | x | ● | ● | ● |
| Antennapedia | RPRC000512 | x | x | x | x | x | x |
| Armadillo | RPRC003585 | ● | ● | ● | ● | ● | ● |
|  | RPRC003617 | ● | ● | ● | ● | ● | ● |
| Cap-n-collar | RPRC011620 | ● | ● | ● | ● | ● | ● |
| Caudal | RPRC000239 | x | ● | ● | ● | ● | ● |
| Cubitus interruptus | RPRC007162 | ● | ● | ● | ● | ● | ● |
| Decapentaplegic | RPRC000401 | ● | ● | ● | ● | ● | ● |
| Deformed | RPRC000437 | x | x | x | x | x | ● |
| Engrailed | RPRC003110 | x | x | x | x | x | ● |
|  | RPRC000282 | x | ● | x | x | x | ● |
| Fused | RPRC003744 | ● | ● | ● | ● | ● | ● |
| Giant | RPRC001027 | ● | ● | x | ● | ● | ● |
| Gooseberry | RPRC008887 | x | ● | x | ● | x | ● |
| Hairy | RPRC000496 | x | ● | ● | ● | ● | ● |
| Hedgehog | RPRC012384 | ● | ● | ● | ● | ● | ● |
| Huckebein | RPRC007615 | ● | ● | x | ● | ● | ● |
| Hunchback | RPRC000230 | ● | ● | x | ● | ● | ● |
| Knirps | RPRC003216 | x | ● | ● | ● | x | ● |
| Knot | RPRC004534 | x | ● | x | ● | ● | ● |
| Kruppel | RPRC000102 | x | ● | x | x | ● | ● |
| Labial | RPRC000004 | x | x | x | x | x | x |
| Nanos | RPRC002927 | x | x | ● | x | ● | ● |
| Odd paired | RPRC013047 | x | ● | x | x | x | ● |
| Odd skipped | RPRC011812 | x | ● | ● | ● | ● | ● |
| Pangolin | RPRC010782 | x | ● | x | ● | ● | ● |
| Pipsqueak | RPRC012968 | ● | ● | ● | ● | ● | ● |
| Proboscipedia | RPRC002128 | ● | ● | x | ● | ● | ● |
| Rasp | RPRC014615 | ● | ● | ● | ● | ● | ● |
| Sex comb reduced | RPRC000310 | x | x | x | x | x | x |
|  | RPRC000369 | x | x | x | x | x | x |
| Sloppy paired 1 | RPRC000987 | x | ● | ● | ● | ● | ● |
| Smoothed | RPRC004966 | ● | ● | ● | ● | ● | ● |
| Tailless | RPRC007025 | x | x | x | ● | ● | ● |
| Tenascin major | RPRC005422 | ● | ● | ● | ● | ● | ● |
| Torso-like | RPRC006513 | ● | ● | ● | ● | ● | ● |
| Tout-velu | RPRC000548 | ● | ● | ● | ● | ● | ● |
| Ultrabithorax | RPRC000565 | ● | ● | ● | x | x | ● |
| Wingless | RPRC005904 | ● | ● | ● | ● | ● | ● |
| Cactus | RPRC017349 | ● | ● | ● | ● | ● | ● |
| Capicua | RPRC001090 | ● | ● | ● | ● | ● | ● |
| Cappuccino | RPRC003459 | ● | ● | ● | ● | ● | ● |
| Cornichon | RPRC014581 | ● | ● | ● | ● | ● | ● |
| Dorsal | RPRC003790 | ● | ● | ● | ● | ● | ● |
| Easter | RPRC003090 | ● | ● | ● | ● | ● | ● |
| Gastrulation defective | RPRC001191 | ● | ● | ● | ● | ● | ● |
| Nudel | RPRC000049 | x | ● | ● | ● | ● | ● |
| Orb | RPRC001021 | ● | ● | ● | ● | ● | ● |
| Pipe | RPRC005278 | ● | x | x | x | x | ● |
| Rhomboid | RPRC008474 | x | ● | x | ● | ● | ● |
| Spätzle | RPRC002634 | ● | ● | ● | ● | ● | ● |
| Spire | RPRC001974 | ● | ● | ● | ● | ● | ● |
|  | RPRC001975 | ● | ● | ● | ● | ● | ● |
| Squid | RPRC007924 | ● | ● | ● | ● | ● | ● |
| Toll | RPRC009262 | ● | ● | ● | ● | ● | ● |
| Windbeutel | RPRC007848 | ● | ● | ● | ● | ● | ● |
| Armitage | RPRC000215 | x | x | • | • | x | • |
| Aubergine | RPRC013054 | • | • | • | • | • | • |
| Bazooka | RPRC001227 | • | • | • | • | • | • |
| Bicaudal C | RPRC001612 | • | • | • | • | • | • |
|  | RPRC001613 | • | • | • | • | • | • |
| Bicaudal D | RPRC000632 | • | • | • | • | • | • |
|  | RPRC000704 | • | • | • | • | • | • |
|  | RPRC004076 | • | • | • | • | • | • |
| Bruno | RPRC007498 | • | • | • | • | • | • |
| Egalitarian | RPRC012064 | • | • | • | • | • | • |
| Egghead | RPRC011094 | • | • | • | • | • | • |
| Eukaryotic translation initiation factor 4E-1 | RPRC013852 | • | • | • | • | • | • |
| Eukaryotic translation initiation factor 4E-2 | RPRC011976 | • | • | x | • | • | • |
| Eukaryotic translation initiation factor 4E-HP | RPRC009699 | • | • | • | • | • | • |
| Exuperantia | RPRC001415 | • | • | • | • | • | • |
| Germ cell-less | RPRC010622 | • | • | • | • | • | • |
| Licorne | RPRC014787 | • | • | • | • | • | • |
| Lost | RPRC014809 | • | • | • | • | • | • |
| Maelstron | RPRC000606 | • | • | • | • | • | • |
| Mago nashi | RPRC010985 | • | • | • | • | • | • |
| Maternal expression at 31B | RPRC009336 | • | • | • | • | • | • |
| Ovo | RPRC002781 | • | • | • | • | • | • |
| Piwi1 | RPRC000252 | x | x | x | x | x | • |
| Piwi2 | RPRC002460 | • | • | • | • | • | • |
| Piwi3 | RPRC001891 | • | • | • | • | • | • |
| Pointed | RPRC011695 | • | • | • | • | • | • |
| Pumilio | RPRC000720 | • | • | • | • | • | • |
| Rump | RPRC014343 | • | • | • | • | • | • |
| Rumpelstiltskin | RPRC014343 | • | • | • | • | • | • |
| Singed | RPRC006015 | • | • | • | • | • | • |
| Staufen | RPRC010019 | • | • | • | • | • | • |
| Tudor | RPRC012896 | • | • | • | • | • | • |
| Vasa | RPRC009661 | • | • | • | • | • | • |
|  | RPRC009663 | • | • | • | • | • | • |
| Wunen | RPRC004485 | • | • | • | • | • | • |
| Wunen2 | RPRC009835 | • | • | • | • | • | • |

The approach identified 81 expressed genes conserved in *R. prolixus*. We cannot rule out that the absence of a transcript is due to an incomplete transcriptome coverage rather than evolutionary divergence. It is plausible to consider that some genes are expressed at very low levels or in a small subset of cells in the developmental stages analyzed and that enrichment analysis may be necessary for their identification. The genes *gurken* (*grk*), *bicoid* (*bcd*), *oskar* (*osk*), were not included in this searched because they have been reported absent in triatomines ^14^. The annotation in our work showed limitations in the current version of the *R. prolixus* genome as more than one gene ID was identified to correspond to the same assembled transcription unit, which was verified by manual assembly. This implies that the genome needs extensive resequencing although a remarkable progress in the annotation of early genes was put forward by Coelho, et al. ^54^ correctly mapping thousands of transcripts even in non-annotated genomic regions.

It would be expected that genes related to oogenesis are not expressed in eggs at times related to gastrulation and germ band formation (48 *hPL*), as well as zygotic expression genes were not expressed in unfertilized eggs. However, a comparison of the different times of development revealed the identification of a stable number of genes of interest in all the times, not allowing to discriminate between presence or absence of gene expression according to a specific time of development. However, all genes showed a maternal expression. These results agree with the assumed role of the maternal genome as director of virtually all aspects of early animal development ^15,80-83^. Specifically, in *D. melanogaster* and *T. castaneum* a high percentage of maternal transcripts were deposited during oogenesis, and so were detected (approximately 58 – 65 %) in the course of the first hours of embryo development, while zygotic transcription was not detected ^15,25,27,83^. These maternal mRNAs have been reported participating in several biological functions, including the establishment of morphogen gradients ^84-86^, segregation of cell-fate determinants ^87-95^, and targeting of protein synthesis to specialized organelles or cellular domains ^96-100^.

To gain insight into this maternal contribution, ovary, unfertilized and 0 *hPL* transcriptomes were used to examine the expression of *R. prolixus* orthologues of reported maternal genes in *D. melanogaster*. The dataset was comprised of 10.277 sequences, of which the 54 % had a *R. prolixus* ortholog to the *D. melanogaster* genes (Figure 3 and Additional file 7). 40,7 % were expressed during the three stages, oogenesis, unfertilized and 0 *hPL* eggs. 1,46 % of the maternal genes had *R. prolixus* orthologs which were expressed only during oogenesis but not in deposited eggs, 1,72 % had *R. prolixus* orthologs that were only maternally loaded into unfertilized eggs, and 2,03 % showed only expression in the first hours of fertilized eggs. At difference to the maternal genes reported in *T. castaneum*, here we identified 306 maternal orthologs more to *D. melanogaster* ^15^. Out of 81 developmental genes investigated, eight were not reported to be maternal in both *D. melanogaster* and *T. castaneum* (Table 4). These eight genes were found in unfertilized eggs of *R. prolixus* (*gooseberry:* RPRC008887, *odd paired*: RPRC013047, *odd skipped*: RPRC011812, *rhomboid:* RPRC008474, *sloppy paired:* RPRC000987) or in both, vitellogenic ovary and unfertilized eggs transcriptomes (*windbeutel*: RPRC007848 and *proboscipedia:* RPRC002128). We also identified a subset of transcript derived from the *Hox* cluster *abdominal-A* (RPRC000598), *Abdominal-B* (RPRC012974), *Deformed* (RPRC000437), *proboscipedia* and *Ultrabithorax* (RPRC000565) (Table 3A, Table 4), suggesting differential maternal/zygotic regulation of the HOX genes in *R. prolixus* that deserves further investigation and that are out of the scope of this work ^101^.

**Figure 3.**
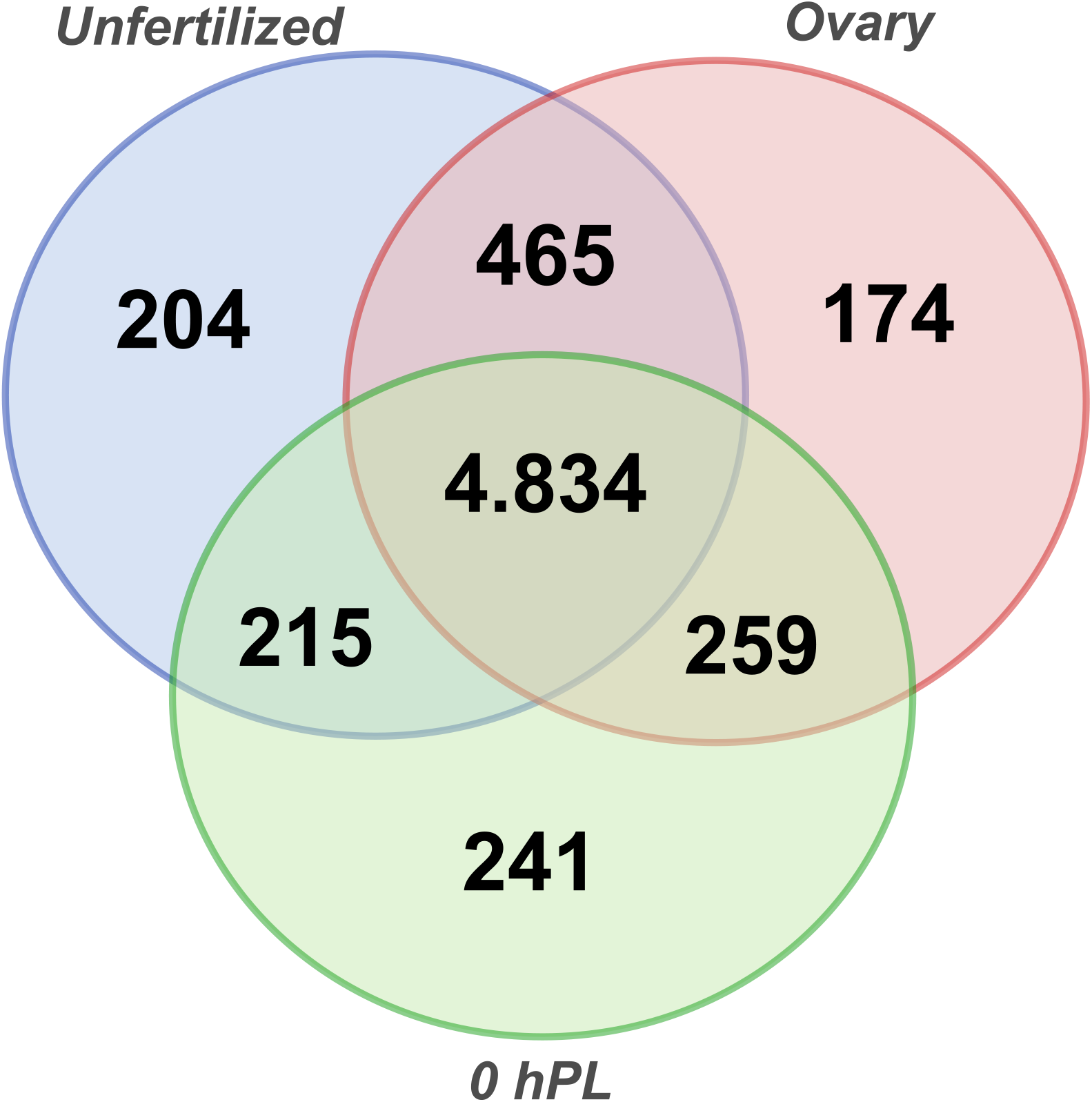
Venn diagram depicting the maternal transcripts across developmental stages. Comparison among the expressed transcripts of the Vitellogenic ovary, Unfertilized and 0 *hPL*, eggs and reference maternal genes.

**Table 4.**
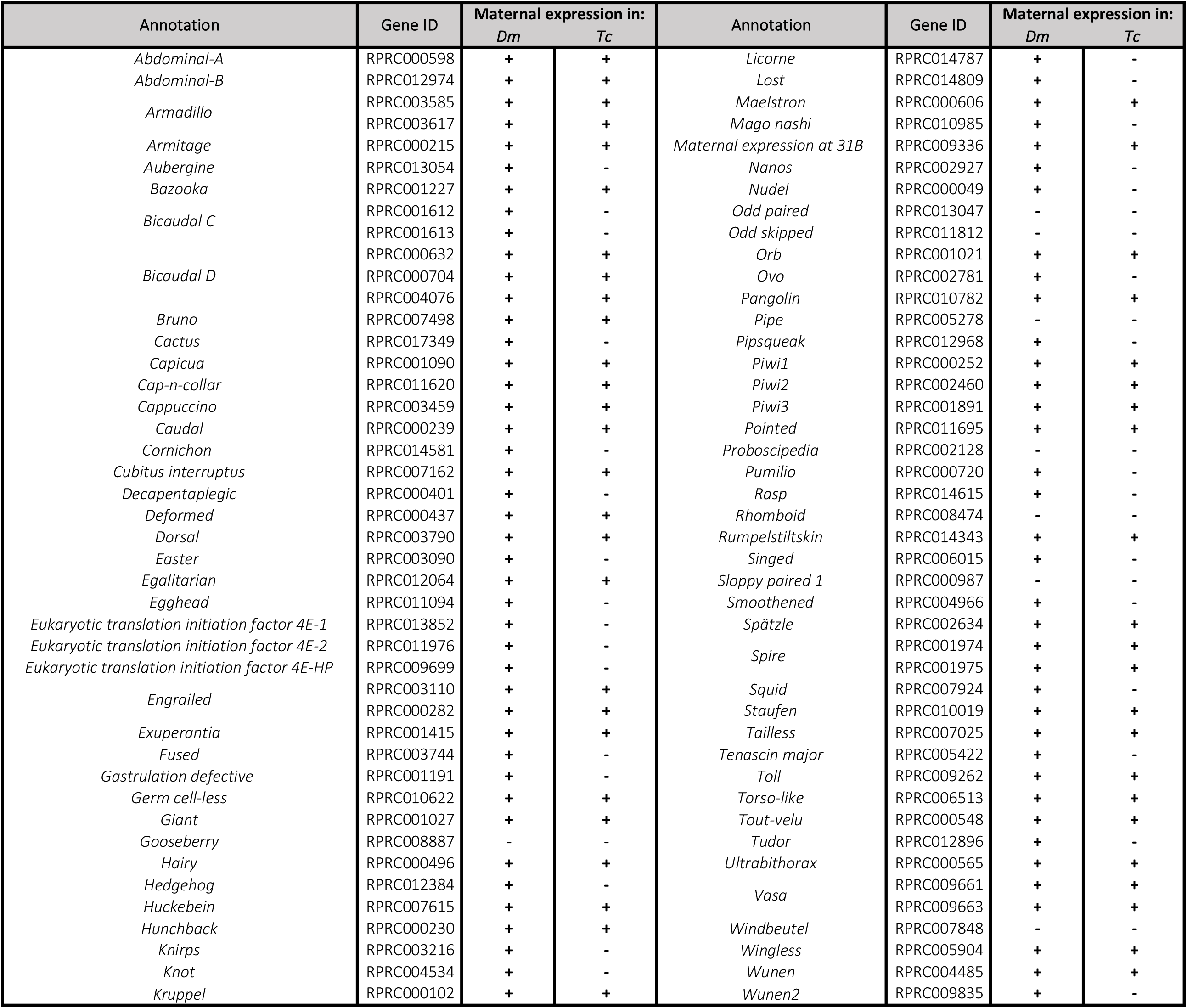
Annotated developmental genes identified as maternal gene in *R. prolixus*. +: detected with maternal expression; -: not detected.

Taken together, our data support the notion that maternal expression of developmental genes that are mostly zygotic in other species might be widespread in *R. prolixus* and the idea that the expression of maternal genes is maintained (either by stability or zygotic expression) during early developmental stages. The results agree with other gene specific studies that have reported maternal expression in the triatomine, during oogenesis and embryo development ^46,48,50,69^.

### 3. Maternal expression validation

To confirm the gene expression patterns revealed by the transcriptomic analyses, 11 genes (from the 81 previously annotated) involved in oogenesis and early embryo development were chosen for qRT-PCR. This analysis was performed with three independent biological replicates different from those used for RNA-seq analysis. The relative expression levels of these genes over time are shown in Figure 3. All of the selected genes showed expression patterns consistent with the results derived from the transcriptome analysis, indicating that our approach, although not intended to be quantitative, was valid for the identification of expressed genes. All genes analyzed showed expression in unfertilized eggs, which confirm that the mRNAs were maternally provided. The different expression level between the vitellogenic ovary and the unfertilized egg could be explained by the nature of the samples, which implied ovarioles depleted of choriogenic egg chambers. *Rp-sqd* showed greater relative expression level than the other genes of interest at the different analyzed times.

### 4. Expression and functional analysis of *Bicaudal D* (*Rp-BicD*) during oogenesis and embryonic development

To further extend the analysis, *in situ* hybridization was performed to evaluated if *Rp-BicD* transcript (Additional file 8) display stage- or tissue-specific expression patterns during oogenesis and/or early development of *R. prolixus*. The structure of an ovariole is shown in Figure 4A-C. The expression of *Rp-BicD* is cytoplasmic and the mRNA is detected in both, the germarium and follicular epithelium of previtellogenic and vitellogenic oocytes. In the germarium, *Rp-BicD* expression is enriched in the region close to the vitellarium (Z3; Figure 4D-E). The expression of *Rp-BicD* is also detected in unfertilized eggs confined in the central region of the egg, with a diffusion of the signal towards the embryo surface (Figure 4F). In embryos up to the onset of blastoderm stage, *Rp-BicD* mRNA could not be detected (data not shown), in agreement with our qRT-PCR data, although we cannot rule out transcripts below the detection limit of the *in situ* hybridization technique. It also shows similarity with *D. melanogaster*, the only insect species in which *BicD* was studied -*BicD* expression drops just prior to cellular blastoderm formation ^102^. This dynamic of expression is similar to that be observed in other maternally supplied RNAs, in which the maternal transcripts persist during the very early embryonic cleavage stages but are degraded just prior to the reach the stage of blastoderm cellularization ^9,103^.

**Figure 4.**
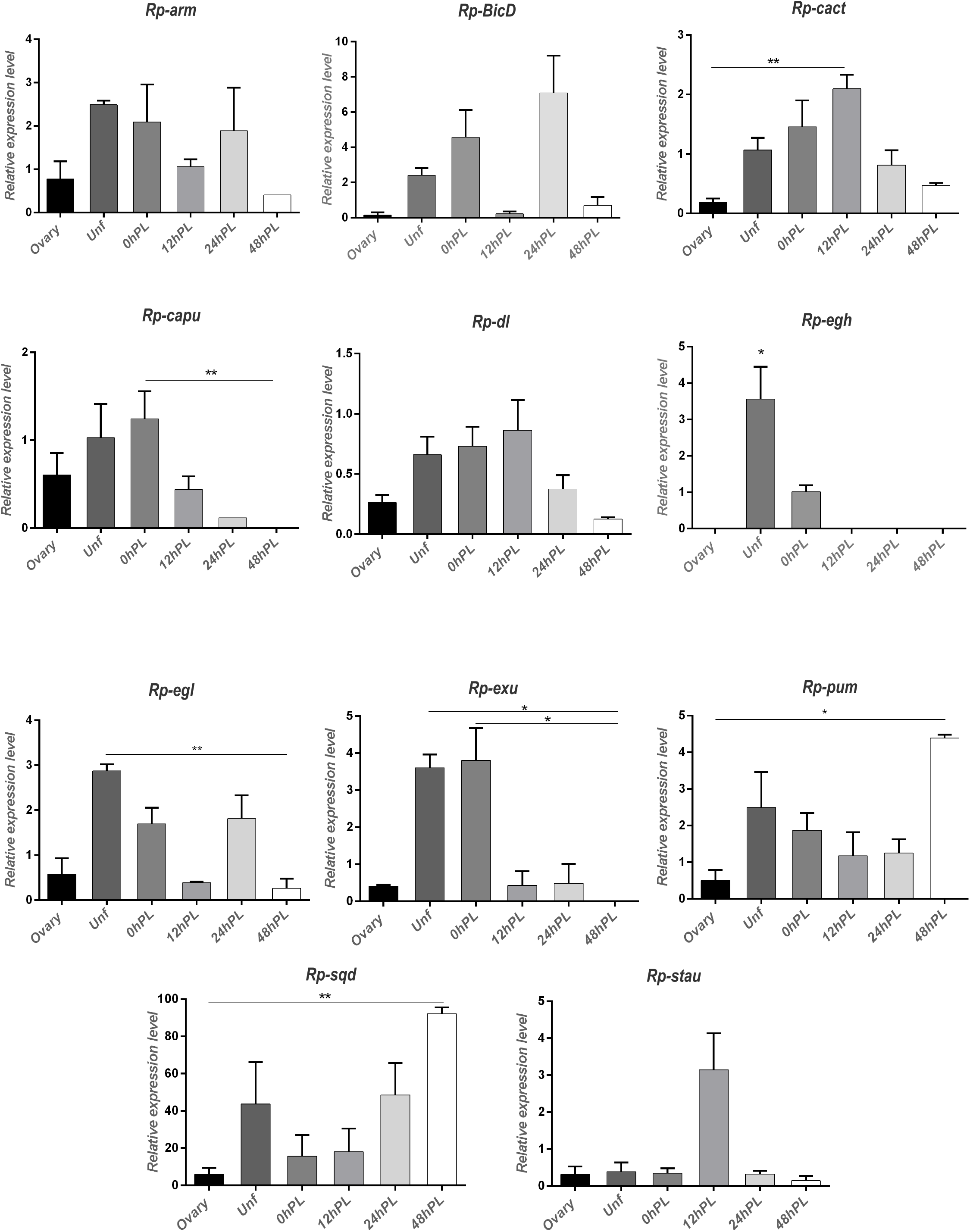
Real-time quantitative PCR of expression of candidate genes in the different stages. X axis: developmental times analyzed. Y axis: expression relative to the reference gene. Values are expressed as mean ± SEM of 3 independent experiments. *Rp-arm*: *Armadillo; Rp-BicD: Bicaudal D; Rp-cact: Cactus; Rp-capu: Cappuccino; Rp-dl: Dorsal; Rp-egh: Egghead; Rp-egl: Egalitarian; Rp-exu: Exuperantia; Rp-pum: Pumilio; Rp-sqd: Squid; Rp-stau: Staufen*. Graphs were performed using GraphPad Prism 7. *<0,1; **<0,05.

**Figure 5.**
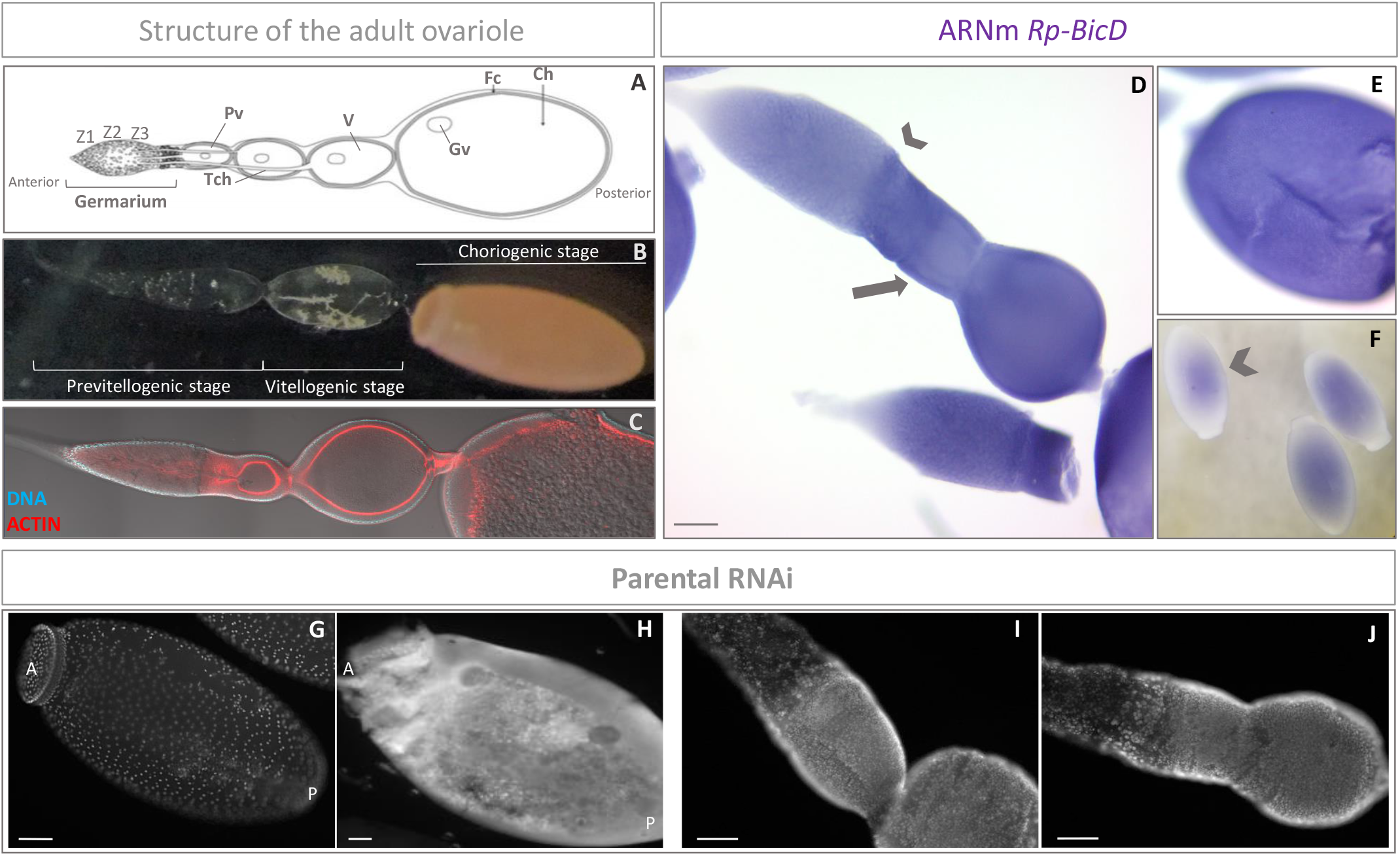
Silencing of *Rp-BicD* produces anembryonic eggs. **A** Schematic of the ovariole showing the germarium host mitotically active cells (i.e. nurse cells) in zone 1 (Z1), zone 2 (Z2) and zone 3 (Z3), and previtellogenic (Pv), vitellogenic (V) and choriogenic (Ch) oocytes. Each oocyte becomes encapsulated by follicle cells (Fc), and remains connected to the germarium through the trophic cords (Tch). Germinal vesicle (Gv). **B** Structure of the ovariole showing the different stages that characterized oogenesis: previtellogenic, vitellogenic and choriogenic stage. **C** Ovariole of a control female showing the nuclei distribution by DAPI staining and the actin filaments by phalloidin staining ^50,116^. **D** Detection of *Rp-BicD* transcript in the ovarioles by *in situ* hybridization. The arrowhead indicates the expression in Z3 of the germarium, the arrow indicates the expression in the vitellogenic oocyte. Scale Bar: 100 µm. **E** Different focal plane of the vitellogenic oocyte showed in **D**. Note the expression of *Rp-BicD* in the follicular cells. **F** Detection of *Rp-BicD* transcript in unfertilized eggs by *in situ* hybridization. The arrowhead indicated the pattern of expression. **G** Egg from control females showing nuclei distribution by DAPI staining. Scale bar: 200 µm. A: Anterior pole of the egg. P: Posterior pole of the egg. **H** Egg from silenced (RNAi^*BicD*^) females showing nuclei distribution by DAPI staining. Scale bar: 100 µm. A: Anterior pole of the egg. P: Posterior pole of the egg. The development stages corresponding to both, control (G) and silenced (H), represent a germ band extension. **I** Ovariole of a control females showing the nuclei distribution by DAPI staining. Scale bar: 100 µm. **J** Ovariole of a silenced (RNAi^*BicD*^) females showing the nuclei distribution by DAPI staining. Scale bar: 100 µm.

In order to determine the function of *Rp-BicD* we performed parental RNAi. We injected non-fed virgin females (n=20) with different concentrations (0.5 to 2.5 µg/female) of dsRNA^*BicD*^. As control, we used dsRNA dsRNA^*β- lac*^ (see methods). After feeding and mating, dsRNA^*BicD*^ and dsRNA^*β- lac*^ injected females were evaluated for survival, egg deposition, embryo lethality, and ovary and embryo phenotype (Additional file 9). In our hands, 45% females injected with dsRNA^*BicD*^ survived until the 5th day after injection and did not reach the oviposition phase. From the survivors (n=9), we investigated the number of egg laid and incubated them to develop for the expected time of embryogenesis to finish (>14 days) to determine lethality. Compared to the control ones, the silenced females laid less eggs (n=43), and of these only 9%, corresponding to the cohort from females injected with the lowest concentration of dsRNA^*BicD*^, hatched to first-instar larvae. The dissection of the not hatching eggs did not show any distinguishable embryonic structure, suggesting that *BicD* might act at very early stages of embryogenesis (Figure 4G-H). The morphology of the ovary analyzed under the dissection microscope and the cellular pattern, as judged by nuclear staining, did not show conspicuous differences. Therefore, the silencing of *Rp*-*BicD* did not alter the normal morphology of the ovaries (Figure 4I-J). The results are similar to the *BicD* phenotypes of *D. melanogaster*. In heterozygous *BicD* females the progression into oogenesis is not affected, but causes sterility or inviable embryos that consist only of a mirror-image duplication of 2–4 posterior segments ^104,105^. Homozygous *BicD* females show ovaries with no diploid germ cell nuclei visible in older egg chambers and no oocyte develops. Also the lack of *BicD* has effect in the zygotic viability, in which null flies die as pupae or young adults ^106^. In *D. melanogaster, BicD* and *Egalitarian* (*Egl*) are part of a complex that transports and localize mRNAs in the early oocyte and in the early embryos, crucial for specifying their respective anteroposterior and dorsoventral axes ^104,107-109^. *Egl* and *BicD* homologues have been identified in *Caenorhabditis elegans* and mammals, and proposed as a part of an evolutionarily conserved cytoskeletal system for transporting transcripts and others ^110,111^. Our qRT-PCR results show similar expression dynamics of *Rp-BicD* and *Rp-egl*, suggesting that these co-expressed trajectories might be associated to establishment of *BicD*/*Egl* localization machinery in *R. prolixus*. Although genes involved in this polarization are well known in *D. melanogaster*, but less so in other insects, include *R. prolixu*s, our results provide evidence that *Rp-BicD* might be an upstream gene in a cascade that contributes to set the axis and the embryonic patterning.

## Conclusions

Our data provide a framework for a better understanding of *R. prolixus* embryogenesis, and complement the remarkable recent work from Coelho, et al. ^54^ that contributed to a better annotation of the *R. prolixus* genome and provided a deep insight into the maternal genes complement of this hemimetabolous insect. In addition, several recent studies showed the expression and phenotype of different genes with maternal contribution. Based on the available data, we propose at least three hierarchies of maternal genes in *R. prolixus*:

1. <u>Genes affecting the early oogenesis</u>, that results in germ cell survival and/or atresic ovaries: *Rp-ATG-8* ^112^; *Rp-cactus* ^46^; *Rp-Piwi-2, Rp-Piwi-3, Rp-Argonauta-3* ^48^; *Rp-Me31B* (Pascual and Rivera Pomar, unpublished data)
2. <u>Gene affecting late oogenesis</u>, that results in eggs with less o incomplete yolk load or impaired choriogenesis: *Rp-Bicaudal C* ^50^; *ULK/Rp-ATG-1* ^113^; *Rp-ATG-3* ^114^; *Rp-ATG-6*^*115*^.
3. <u>Genes affecting embryonic patterning</u>, that results in embryo lethality: *Rp-giant* ^69^; *Rp-dorsal* ^46^; *Rp-Bicaudal D* (this work),

Genome analysis opened a broad and exciting path to study oogenesis and to decipher the genetic pathways of oogenesis in this emerging model organism, adding to the already well known cellular processes established by the pioneer work of Erwin Huebner. These studies will enrich the developmental evolutionary studies in the context of insect comparative genomics, and, in the case of *R. prolixus*, could contribute to devise new strategies to control Chagas disease insects.

## Supporting information

Suppelentary 1B

## Acknowledgements

The authors thank all members of Rivera-Pomar lab, in particular to Elías Gazza and Agustín Rolandelli for fruitful discussions, and to Viviana Decker (Instituto Nacional de Tecnología Agropecuaria, Pergamino) for kindly share the qRT-PCR facility. R.R-P is professor at UNNOBA and investigator of CONICET. A.P is postdoctoral fellow of CONICET and UNNOBA. This work was funded by ANPCyT (PICT-2013-1554 to RR-P), UNNOBA (SIB 0626/2019 to RR-P), and Alexander von Humboldt Research Prize to RR-P.

## Supporting information (Excel files available under request)

**Additional file 1:** Maternal *D. melanogaster* genes, reported as FlyBase symbol.

**Additional file 2:** Primers used by RT-qPCR assays and *Rp-BicD* experiments.

**Additional file 3:** COG functional category annotation using eggNOG-mapper.

**Additional file 4:** InterproScan results.

**Additional file 5:** GO-term enrichment.

**Additional file 6:** Go-term enrichment form annotated transcripts common to all six developmental stages.

**Additional file 7:** BLASTX from maternal genes.

**Additional file 8:** The predicted sequences were aligned with orthologues from other species with Clustal Ω ^117^. Alignment of the protein sequence of *Rp-BicD* with orthologues species, extracted from NCBI sequence database *D. melanogaster* (NP_001260531.1), *Tribolium castaneum* (EFA07458.1), *Culex quinquefasciatus* (EDS37197.1), *Anopheles darlingi* (ETN62092.1), *Daphnia magna* JAN71328.1, *Homo sapiens* (AAB94805.1), *Mus musculus* (NP_001034268.1), *Caenorhabditis elegans* (CDK13419.1). The amino acid conservation is visualized with black blocks, the amino acid group level conservation with gray blocks.

**Additional file 9:** Summary of Parental RNAi experiment.

## References

1 Moussian, B. & Roth, S. Dorsoventral axis formation in the Drosophila embryo--shaping and transducing a morphogen gradient. Current biology : CB 15, R887–899, doi:10.1016/j.cub.2005.10.026 (2005).

2 Dzamba, B. J. & DeSimone, D. W. Extracellular Matrix (ECM) and the Sculpting of Embryonic Tissues. Curr Top Dev Biol 130, 245–274, doi:10.1016/bs.ctdb.2018.03.006 (2018).

3 Salazar-Ciudad, I. Morphological evolution and embryonic developmental diversity in metazoa. Development 137, 531–539, doi:10.1242/dev.045229 (2010).

4 Davis, G. K. & Patel, N. H. Short, long, and beyond: molecular and embryological approaches to insect segmentation. Annual review of entomology 47, 669–699, doi:10.1146/annurev.ento.47.091201.145251 (2002).

5 Liu, P. Z. & Kaufman, T. C. Short and long germ segmentation: unanswered questions in the evolution of a developmental mode. Evolution & development 7, 629–646, doi:10.1111/j.1525-142X.2005.05066.x (2005).

6 El-Sherif, E., Lynch, J. A. & Brown, S. J. Comparisons of the embryonic development of Drosophila, Nasonia, and Tribolium. Wiley interdisciplinary reviews. Developmental biology 1, 16–39, doi:10.1002/wdev.3 (2012).

7 Mellanby, H. Memoirs: The Early Embryonic Development of Rhodnius Prolixus (Hemiptera, Heteroptera). Quarterly Journal of Microscopical Science s2-78, 71–90 (1935).

8 Lynch, J. A. & Roth, S. The evolution of dorsal-ventral patterning mechanisms in insects. Genes Dev. 25, 107–118, doi:10.1101/gad.2010711 (2011).

9 Tadros, W. & Lipshitz, H. D. The maternal-to-zygotic transition: a play in two acts. Development 136, 3033–3042, doi:10.1242/dev.033183 (2009).

10 Behura, S. K. et al. Comparative genomic analysis of Drosophila melanogaster and vector mosquito developmental genes. PloS one 6, e21504, doi:10.1371/journal.pone.0021504 (2011).

11 Harker, B. W. et al. Stage-specific transcription during development of Aedes aegypti. BMC developmental biology 13, 29, doi:10.1186/1471-213X-13-29 (2013).

12 Ewen-Campen, B. et al. The maternal and early embryonic transcriptome of the milkweed bug Oncopeltus fasciatus. BMC genomics 12, 61, doi:10.1186/1471-2164-12-61 (2011).

13 Fan, X. B. et al. An Overview of Embryogenesis: External Morphology and Transcriptome Profiling in the Hemipteran Insect Nilaparvata lugens. Frontiers in physiology 11, 106, doi:10.3389/fphys.2020.00106 (2020).

14 Lavore, A. et al. Comparative analysis of zygotic developmental genes in Rhodnius prolixus genome shows conserved features on the tracheal developmental pathway. Insect Biochem. Mol. Biol. 64, 32–43, doi:10.1016/j.ibmb.2015.06.012 (2015).

15 Preuss, K. M., Lopez, J. A., Colbourne, J. K. & Wade, M. J. Identification of maternally-loaded RNA transcripts in unfertilized eggs of Tribolium castaneum. BMC Genomics 13, 671, doi:10.1186/1471-2164-13-671 (2012).

16 Hu, X., Ke, L., Wang, Z. & Zeng, Z. Dynamic transcriptome landscape of Asian domestic honeybee (Apis cerana) embryonic development revealed by high-quality RNA sequencing. BMC developmental biology 18, 11, doi:10.1186/s12861-018-0169-1 (2018).

17 Simon, S. et al. Comparative transcriptomics reveal developmental turning points during embryogenesis of a hemimetabolous insect, the damselfly Ischnura elegans. Scientific reports 7, 13547, doi:10.1038/s41598-017-13176-8 (2017).

18 Clark, E. Dynamic patterning by the Drosophila pair-rule network reconciles long-germ and short-germ segmentation. PLoS biology 15, e2002439, doi:10.1371/journal.pbio.2002439 (2017).

19 Clark, E. & Peel, A. D. Evidence for the temporal regulation of insect segmentation by a conserved sequence of transcription factors. Development, doi:10.1242/dev.155580 (2018).

20 Damen, W. G., Janssen, R. & Prpic, N. M. Pair rule gene orthologs in spider segmentation. Evolution & development 7, 618–628, doi:10.1111/j.1525-142X.2005.05065.x (2005).

21 Farzana, L. & Brown, S. J. Hedgehog signaling pathway function conserved in Tribolium segmentation. Development genes and evolution 218, 181–192, doi:10.1007/s00427-008-0207-2 (2008).

22 Janssen, R. & Budd, G. E. Deciphering the onychophoran ‘segmentation gene cascade’: Gene expression reveals limited involvement of pair rule gene orthologs in segmentation, but a highly conserved segment polarity gene network. Developmental biology 382, 224–234, doi:10.1016/j.ydbio.2013.07.010 (2013).

23 Benton, M. A. A revised understanding of Tribolium morphogenesis further reconciles short and long germ development. PLoS Biol. 16, e2005093, doi:10.1371/journal.pbio.2005093 (2018).

24 Oppenheim, S. J., Baker, R. H., Simon, S. & DeSalle, R. We can’t all be supermodels: the value of comparative transcriptomics to the study of non-model insects. Insect molecular biology 24, 139–154, doi:10.1111/imb.12154 (2015).

25 Arbeitman, M. N. et al. Gene expression during the life cycle of Drosophila melanogaster. Science 297, 2270–2275, doi:10.1126/science.1072152 (2002).

26 De Renzis, S., Elemento, O., Tavazoie, S. & Wieschaus, E. Unmasking activation of the zygotic genome using chromosomal deletions in the Drosophila embryo. PloS one 5, 1036–1051 (2007).

27 Lecuyer, E. et al. Global analysis of mRNA localization reveals a prominent role in organizing cellular architecture and function. Cell 131, 174–187, doi:10.1016/j.cell.2007.08.003 (2007).

28 Tomancak, P. et al. Global analysis of patterns of gene expression during Drosophila embryogenesis. Genome biology 8, R145, doi:10.1186/gb-2007-8-7-r145 (2007).

29 Tomancak, P. et al. Systematic determination of patterns of gene expression during Drosophila embryogenesis. Genome biology 3, RESEARCH0088, doi:10.1186/gb-2002-3-12-research0088 (2002).

30 Pridohl, F. et al. Transcriptome sequencing reveals maelstrom as a novel target gene of the terminal system in the red flour beetle Tribolium castaneum. Development 144, 1339–1349, doi:10.1242/dev.136853 (2017).

31 Amsel, D., Vilcinskas, A. & Billion, A. Evaluation of high-throughput isomiR identification tools: illuminating the early isomiRome of Tribolium castaneum. BMC Bioinformatics 18, 359, doi:10.1186/s12859-017-1772-z (2017).

32 Coura, J. R. & Borges-Pereira, J. Chagas disease. What is known and what should be improved: a systemic review. Rev Soc Bras Med Trop 45, 286–296, doi:10.1590/s0037-86822012000300002 (2012).

33 Chagas, C. R. J. Nova tripanosomíase humana. Estudos sobre a morphologia e o ciclo evolutivo do Schizotrypanum cruzi n. gen. n. esp., agente da nova entidade mórbida do homem. . Mem Inst Oswaldo Cruz. 1, 159–218. (1909).

34 Wigglesworth, V. B. The hormonal regularion of growth and reproduction in insects. Adv. Insect Physiol 2 (1964).

35 Wigglesworth, V. B. The origin of sensory neurons in an insect. Q. J. Microscopy Sci 93, 93–112 (1953).

36 Wigglesworth, V. B. The physiology of ecdysis in Rhodnius prolixus (Hemiptera). II. Factors controlling moulting and metamorphosis. Q. J. Microscopy Sci 77, 191–222 (1934).

37 Wigglesworth, V. B. The Principles of Insect Physiology. Methuen, London (1939).

38 Huebner, E. & Anderson, E. A cytological study of the ovary of Rhodnius prolixus. Cytoarchitecture and development of the trophic chamber. J. Morphol. 138, 1–40, doi:10.1002/jmor.1051380102 (1972).

39 Huebner, E. & Anderson, E. A cytological study of the ovary of Rhodnius prolixus. I. The ontogeny of the follicular epithelium. J. Morphol. 136, 459–493, doi:10.1002/jmor.1051360405 (1972).

40 Huebner, E. & Anderson, E. A cytological study of the ovary of Rhodnius prolixus. II. Oocyte differentiation. J. Morphol. 137, 385–415, doi:10.1002/jmor.1051370402 (1972).

41 Lutz, D. A. & Huebner, E. Development and cellular differentiation of an insect telotrophic ovary (Rhodnius prolixus). Tissue Cell 12, 773–794, doi:10.1016/0040-8166(80)90029-4 (1980).

42 Lutz, D. A. & Huebner, E. Development of nurse cell-oocyte interactions in the insect telotrophic ovary (Rhodnius prolixus). Tissue Cell 13, 321–335, doi:10.1016/0040-8166(81)90008-2 (1981).

43 Huebner, E. Nurse cell-oocyte interaction in the telotrophic ovarioles of an insect, Rhodnius prolixus. Tissue Cell 13, 105–125, doi:10.1016/0040-8166(81)90042-2 (1981).

44 Huebner, E. Oocyte-follicle cell interaction during normal oogenesis and atresia in an insect. J Ultrastruct Res 74, 95–104, doi:10.1016/s0022-5320(81)80112-8 (1981).

45 Mesquita, R. D. et al. Genome of Rhodnius prolixus, an insect vector of Chagas disease, reveals unique adaptations to hematophagy and parasite infection. Proc Natl Acad Sci U S A 112, 14936–14941, doi:10.1073/pnas.1506226112 (2015).

46 Berni, M. et al. Toll signals regulate dorsal-ventral patterning and anterior-posterior placement of the embryo in the hemipteran Rhodnius prolixus. Evodevo 5, 38, doi:10.1186/2041-9139-5-38 (2014).

47 Lavore, A., Esponda-Behrens, N., Pagola, L. & Rivera-Pomar, R. The gap gene Kruppel of Rhodnius prolixus is required for segmentation and for repression of the homeotic gene sex comb-reduced. Dev. Biol. 387, 121–129, doi:10.1016/j.ydbio.2013.12.030 (2014).

48 Brito, T. et al. Transcriptomic and functional analyses of the piRNA pathway in the Chagas disease vector Rhodnius prolixus. PLoS neglected tropical diseases 12, e0006760, doi:10.1371/journal.pntd.0006760 (2018).

49 Nunes-da-Fonseca, R., Berni, M., Tobias-Santos, V., Pane, A. & Araujo, H. M. Rhodnius prolixus: From classical physiology to modern developmental biology. Genesis 55, doi:10.1002/dvg.22995 (2017).

50 Pascual, A., Vilardo, E. S., Taibo, C., Sabioy García, J. & Rivera-Pomar, R. Bicaudal C is required for the function of the follicular epithelium during oogenesis in Rhodnius prolixus. Dev. Genes Evol., doi:https://doi.org/10.1007/s00427-021-00673-0 (2021).

51 Leyria, J., Orchard, I. & Lange, A. B. Transcriptomic analysis of regulatory pathways involved in female reproductive physiology of Rhodnius prolixus under different nutritional states. Scientific reports 10, 11431, doi:10.1038/s41598-020-67932-4 (2020).

52 Leyria, J., Orchard, I. & Lange, A. B. What happens after a blood meal? A transcriptome analysis of the main tissues involved in egg production in Rhodnius prolixus, an insect vector of Chagas disease. PLoS neglected tropical diseases 14, e0008516, doi:10.1371/journal.pntd.0008516 (2020).

53 Medeiros, M. N. et al. Transcriptome and gene expression profile of ovarian follicle tissue of the triatomine bug Rhodnius prolixus. Insect Biochem. Mol. Biol. 41, 823–831, doi:10.1016/j.ibmb.2011.06.004 (2011).

54 Coelho, V. L. et al. Analysis of ovarian transcriptomes reveals thousands of novel genes in the insect vector Rhodnius prolixus. Scientific reports 11, 1918, doi:10.1038/s41598-021-81387-1 (2021).

55 Altschul, S. F., Gish, W., Miller, W., Myers, E. W. & Lipman, D. J. Basic local alignment search tool. J. Mol. Biol. 215, 403–410, doi:10.1016/S0022-2836(05)80360-2 (1990).

56 Quast, C. et al. The SILVA ribosomal RNA gene database project: improved data processing and web-based tools. Nucleic acids research 41, D590–596, doi:10.1093/nar/gks1219 (2013).

57 Kim, D. et al. TopHat2: accurate alignment of transcriptomes in the presence of insertions, deletions and gene fusions. Genome biology 14, R36, doi:10.1186/gb-2013-14-4-r36 (2013).

58 Giraldo-Calderon, G. I. et al. VectorBase: an updated bioinformatics resource for invertebrate vectors and other organisms related with human diseases. Nucleic Acids Res. 43, D707–713, doi:10.1093/nar/gku1117 (2015).

59 Wang, L., Wang, S. & Li, W. RSeQC: quality control of RNA-seq experiments. Bioinformatics 28, 2184–2185, doi:10.1093/bioinformatics/bts356 (2012).

60 Garcia-Alcalde, F. et al. Qualimap: evaluating next-generation sequencing alignment data. Bioinformatics 28, 2678–2679, doi:10.1093/bioinformatics/bts503 (2012).

61 Garber, M., Grabherr, M. G., Guttman, M. & Trapnell, C. Computational methods for transcriptome annotation and quantification using RNA-seq. Nat. Methods 8, 469–477, doi:10.1038/nmeth.1613 (2011).

62 Trapnell, C. et al. Differential gene and transcript expression analysis of RNA-seq experiments with TopHat and Cufflinks. Nat Protoc 7, 562–578, doi:10.1038/nprot.2012.016 (2012).

63 Simao, F. A., Waterhouse, R. M., Ioannidis, P., Kriventseva, E. V. & Zdobnov, E. M. BUSCO: assessing genome assembly and annotation completeness with single-copy orthologs. Bioinformatics 31, 3210–3212, doi:10.1093/bioinformatics/btv351 (2015).

64 Seemann, T. & Gladman, S. Fasta Statistics: Display summary statistics for a fasta file, <https://github.com/galaxyproject/tools-iuc> (2012).

65 Haas, B. J. et al. De novo transcript sequence reconstruction from RNA-seq using the Trinity platform for reference generation and analysis. Nat Protoc 8, 1494–1512, doi:10.1038/nprot.2013.084 (2013).

66 Zdobnov, E. M. & Apweiler, R. InterProScan--an integration platform for the signature-recognition methods in InterPro. Bioinformatics 17, 847–848, doi:10.1093/bioinformatics/17.9.847 (2001).

67 topGO: Enrichment Analysis for Gene Ontology v. 2.40.0 (Bioconductor, R package, 2020).

68 Ginzburg, N., Cohen, M. & Chipman, A. D. Factors involved in early polarization of the anterior-posterior axis in the milkweed bug Oncopeltus fasciatus. Genesis 55, doi:10.1002/dvg.23027 (2017).

69 Lavore, A., Pagola, L., Esponda-Behrens, N. & Rivera-Pomar, R. The gap gene giant of Rhodnius prolixus is maternally expressed and required for proper head and abdomen formation. Dev. Biol. 361, 147–155, doi:10.1016/j.ydbio.2011.06.038 (2012).

70 Folmes, C. D. & Terzic, A. Metabolic determinants of embryonic development and stem cell fate. Reproduction, fertility, and development 27, 82–88, doi:10.1071/RD14383 (2014).

71 Miyazawa, H. & Aulehla, A. Revisiting the role of metabolism during development. Development 145, doi:10.1242/dev.131110 (2018).

72 Atella, G. C. et al. Oogenesis and egg development in triatomines: a biochemical approach. An Acad Bras Cienc 77, 405–430, doi:S0001-37652005000300005 (2005).

73 Fraga, A. et al. Glycogen and glucose metabolism are essential for early embryonic development of the red flour beetle Tribolium castaneum. PloS one 8, e65125, doi:10.1371/journal.pone.0065125 (2013).

74 Vital, W. et al. Germ band retraction as a landmark in glucose metabolism during Aedes aegypti embryogenesis. BMC Dev. Biol. 10, 25, doi:10.1186/1471-213X-10-25 (2010).

75 Moraes, J. et al. Glucose metabolism during embryogenesis of the hard tick Boophilus microplus. Comparative biochemistry and physiology. Part A, Molecular & integrative physiology 146, 528–533, doi:10.1016/j.cbpa.2006.05.009 (2007).

76 Santos, R. et al. Carbohydrate accumulation and utilization by oocytes of Rhodnius prolixus. Arch. Insect Biochem. Physiol. 67, 55–62, doi:10.1002/arch.20217 (2008).

77 Santos, R., Rosas-Oliveira, R., Saraiva, F. B., Majerowicz, D. & Gondim, K. C. Lipid accumulation and utilization by oocytes and eggs of Rhodnius prolixus. Arch. Insect Biochem. Physiol. 77, 1–16, doi:10.1002/arch.20414 (2011).

78 Pontes, E. G., Leite, P., Majerowicz, D., Atella, G. C. & Gondim, K. C. Dynamics of lipid accumulation by the fat body of Rhodnius prolixus: the involvement of lipophorin binding sites. J. Insect Physiol. 54, 790–797, doi:10.1016/j.jinsphys.2008.02.003 (2008).

79 Fruttero, L. L., Frede, S., Rubiolo, E. R. & Canavoso, L. E. The storage of nutritional resources during vitellogenesis of Panstrongylus megistus (Hemiptera: Reduviidae): the pathways of lipophorin in lipid delivery to developing oocytes. J. Insect Physiol. 57, 475–486, doi:10.1016/j.jinsphys.2011.01.009 (2011).

80 Baugh, L. R., Hill, A. A., Slonim, D. K., Brown, E. L. & Hunter, C. P. Composition and dynamics of the Caenorhabditis elegans early embryonic transcriptome. Development 130, 889–900, doi:10.1242/dev.00302 (2003).

81 Wei, Z., Angerer, R. C. & Angerer, L. M. A database of mRNA expression patterns for the sea urchin embryo. Developmental biology 300, 476–484, doi:10.1016/j.ydbio.2006.08.034 (2006).

82 Wang, Q. T. et al. A genome-wide study of gene activity reveals developmental signaling pathways in the preimplantation mouse embryo. Developmental cell 6, 133–144, doi:10.1016/s1534-5807(03)00404-0 (2004).

83 Lefebvre, F. A. & Lecuyer, E. Flying the RNA Nest: Drosophila Reveals Novel Insights into the Transcriptome Dynamics of Early Development. Journal of developmental biology 6, doi:10.3390/jdb6010005 (2018).

84 Driever, W. & Nusslein-Volhard, C. The bicoid protein determines position in the Drosophila embryo in a concentration-dependent manner. Cell 54, 95–104, doi:10.1016/0092-8674(88)90183-3 (1988).

85 Ephrussi, A., Dickinson, L. K. & Lehmann, R. Oskar organizes the germ plasm and directs localization of the posterior determinant nanos. Cell 66, 37–50, doi:10.1016/0092-8674(91)90137-n (1991).

86 Gavis, E. R. & Lehmann, R. Localization of nanos RNA controls embryonic polarity. Cell 71, 301–313, doi:10.1016/0092-8674(92)90358-j (1992).

87 Broadus, J., Fuerstenberg, S. & Doe, C. Q. Staufen-dependent localization of prospero mRNA contributes to neuroblast daughter-cell fate. Nature 391, 792–795, doi:10.1038/35861 (1998).

88 Gore, A. V. et al. The zebrafish dorsal axis is apparent at the four-cell stage. Nature 438, 1030–1035, doi:10.1038/nature04184 (2005).

89 Takizawa, P. A., Sil, A., Swedlow, J. R., Herskowitz, I. & Vale, R. D. Actin-dependent localization of an RNA encoding a cell-fate determinant in yeast. Nature 389, 90–93, doi:10.1038/38015 (1997).

90 Hughes, J. R., Bullock, S. L. & Ish-Horowicz, D. Inscuteable mRNA localization is dynein-dependent and regulates apicobasal polarity and spindle length in Drosophila neuroblasts. Current biology : CB 14, 1950–1956, doi:10.1016/j.cub.2004.10.022 (2004).

91 Long, R. M. et al. Mating type switching in yeast controlled by asymmetric localization of ASH1 mRNA. Science 277, 383–387, doi:10.1126/science.277.5324.383 (1997).

92 Neuman-Silberberg, F. S. & Schupbach, T. The Drosophila dorsoventral patterning gene gurken produces a dorsally localized RNA and encodes a TGF alpha-like protein. Cell 75, 165–174 (1993).

93 Simmonds, A. J., dosSantos, G., Livne-Bar, I. & Krause, H. M. Apical localization of wingless transcripts is required for wingless signaling. Cell 105, 197–207, doi:10.1016/s0092-8674(01)00311-7 (2001).

94 Zhang, J. et al. The role of maternal VegT in establishing the primary germ layers in Xenopus embryos. Cell 94, 515–524, doi:10.1016/s0092-8674(00)81592-5 (1998).

95 Melton, D. A. Translocation of a localized maternal mRNA to the vegetal pole of Xenopus oocytes. Nature 328, 80–82, doi:10.1038/328080a0 (1987).

96 Adereth, Y., Dammai, V., Kose, N., Li, R. & Hsu, T. RNA-dependent integrin alpha3 protein localization regulated by the Muscleblind-like protein MLP1. Nature cell biology 7, 1240–1247, doi:10.1038/ncb1335 (2005).

97 Lambert, J. D. & Nagy, L. M. Asymmetric inheritance of centrosomally localized mRNAs during embryonic cleavages. Nature 420, 682–686, doi:10.1038/nature01241 (2002).

98 Lawrence, J. B. & Singer, R. H. Intracellular localization of messenger RNAs for cytoskeletal proteins. Cell 45, 407–415, doi:10.1016/0092-8674(86)90326-0 (1986).

99 Mingle, L. A. et al. Localization of all seven messenger RNAs for the actin-polymerization nucleator Arp2/3 complex in the protrusions of fibroblasts. Journal of cell science 118, 2425–2433, doi:10.1242/jcs.02371 (2005).

100 Zhang, H. L. et al. Neurotrophin-induced transport of a beta-actin mRNP complex increases beta-actin levels and stimulates growth cone motility. Neuron 31, 261–275, doi:10.1016/s0896-6273(01)00357-9 (2001).

101 Esponda-Behrens, N. Estudios funcionales comparados de la evolución de la segmentación en insectos. Facultad de Ciencias Exactas, doi:https://doi.org/10.35537/10915/43085 (2014).

102 Suter, B., Romberg, L. M. & Steward, R. Bicaudal-D, a Drosophila gene involved in developmental asymmetry: localized transcript accumulation in ovaries and sequence similarity to myosin heavy chain tail domains. Genes Dev. 3, 1957–1968 (1989).

103 Salz, H. K. et al. The Drosophila female-specific sexdetermination gene. Sex-lethal, has stage-, tissue-, and sex-specific RNAs suggesting multiple modes of regulation. Genes Dev. 3, 708–719 (1989).

104 Suter, B. & Steward, R. Requirement for phosphorylation and localization of the Bicaudal-D protein in Drosophila oocyte differentiation. Cell 67, 917–926, doi:10.1016/0092-8674(91)90365-6 (1991).

105 Mohler, J. & Wieschaus, E. F. Dominant maternal-effect mutations of Drosophila melanogaster causing the production of double-abdomen embryos. Genetics 112, 803–822 (1986).

106 Ran, B., Bopp, R. & Suter, B. Null alleles reveal novel requirements for Bic-D during Drosophila oogenesis and zygotic development. Development 120, 1233–1242 (1994).

107 Bullock, S. L. & Ish-Horowicz, D. Conserved signals and machinery for RNA transport in Drosophila oogenesis and embryogenesis. Nature 414, 611–616, doi:10.1038/414611a (2001).

108 Mach, J. M. & Lehmann, R. An Egalitarian-BicaudalD complex is essential for oocyte specification and axis determination in Drosophila. Genes Dev. 11, 423–435, doi:10.1101/gad.11.4.423 (1997).

109 Vazquez-Pianzola, P. et al. The mRNA transportome of the BicD/Egl transport machinery. RNA biology 14, 73–89, doi:10.1080/15476286.2016.1251542 (2017).

110 Baens, M. & Marynen, P. A human homologue (BICD1) of the Drosophila bicaudal-D gene. Genomics 45, 601–606, doi:10.1006/geno.1997.4971 (1997).

111 Aguirre-Chen, C., Bulow, H. E. & Kaprielian, Z. C. elegans bicd-1, homolog of the Drosophila dynein accessory factor Bicaudal D, regulates the branching of PVD sensory neuron dendrites. Development 138, 507–518, doi:10.1242/dev.060939 (2011).

112 Pereira, J. et al. Silencing of RpATG8 impairs the biogenesis of maternal autophagosomes in vitellogenic oocytes, but does not interrupt follicular atresia in the insect vector Rhodnius prolixus. PLoS neglected tropical diseases 14, e0008012, doi:10.1371/journal.pntd.0008012 (2020).

113 Bomfim, L. & Ramos, I. Deficiency of ULK1/ATG1 in the follicle cells disturbs ER homeostasis and causes defective chorion deposition in the vector Rhodnius prolixus. FASEB J. 34, 13561–13572, doi:10.1096/fj.202001396R (2020).

114 Santos, A. & Ramos, I. ATG3 Is Important for the Chorion Ultrastructure During Oogenesis in the Insect Vector Rhodnius prolixus. Frontiers in physiology 12, 638026, doi:10.3389/fphys.2021.638026 (2021).

115 Vieira, P. H., Bomfim, L., Atella, G. C., Masuda, H. & Ramos, I. Silencing of RpATG6 impaired the yolk accumulation and the biogenesis of the yolk organelles in the insect vector R. prolixus. PLoS neglected tropical diseases 12, e0006507, doi:10.1371/journal.pntd.0006507 (2018).

116 Pascual, A. Genómica del desarrollo embrionario de Rhodnius prolixus. Facultad de Ciencias Exactas, Área Ciencias Biológicas, doi:https://doi.org/10.35537/10915/90493 (2019).

117 Sievers, F. & Higgins, D. G. Clustal Omega, accurate alignment of very large numbers of sequences. Methods in molecular biology 1079, 105–116, doi:10.1007/978-1-62703-646-7_6 (2014).

