## Supplementary 1A for "Dynamics of maternal gene expression in *Rhodnius prolixus*"

| Primer Name | Sequence 5'-3' | Amplicon length (bp) | PCR efficiency (%) |
|---|---|---|---|
| armFw | CATCTCGCTGGCCTCTTATC | 235 | 98 |
| armRv | TGTAGAGCTCCCACCGTACC | | |
| BicDFw | GCCGAAAAACAGATGGAAGA | 254 | 94,8 |
| BicDRv | TCGACATGCTTTGTCTCTGG | | |
| cactFw | GTGCTGGTGCTTGTACGAAA | 154 | 91,43 |
| cactRv | GGAGTCGGACGATACCTCAA | | |
| capuFw | ACAAAGAGGACAAGCCGATG | 171 | 101 |
| capuRv | TCACTTGGTTCTGGAACTGG | | |
| dlFw | GTCATTGTTCTGGTTCAGTAGC | 181 | 90 |
| dlRv | GGCGTTCGGAATGAAATAGC | | |
| eghFw | ACCCTGAAAGGTAGCCCACT | 245 | 93 |
| eghRv | GCTTTGAACATGGCTCCACT | | |
| eglFw | CCAGGTCTACAACGCTATGC | 189 | 88,4 |
| eglRv | TCTACCAGTGTGGGCAGATG | | |
| exuFw | CACGGAAGGAAGGGATCTTA | 238 | 100 |
| exuRv | ACAGGCATCGTGGCTCTTAT | | |
| pumFw | CCAGCGTATCCTGGAACATT | 220 | 95 |
| pumRv | CTCAACGACGTTTGAAGCAA | | |
| sqdFw | AATTACTTCGCGCAATACGG | 193 | 86 |
| sqdRv | TGCCATCAGGTTTTGGTGTA | | |
| satuFw | AATGGAAATTCACCCACAGC | 186 | 97,4 |
| StauRv | AGTGGCCAGTGATACCAAGC | | |
| αtubFw | CAAATAATTACGCCCGAGGA | 230 | 84,9 |
| αtubRv | TTGAGGAGCTGGGTAAATGG | | |

The primer sequences for dl and cact are from Rolandelli, A. et al., 2020
