## Supplementary material for "Dynamics of maternal gene expression in *Rhodnius prolixus*": Suppelentary 1B

| Name | Sequence 5'-3' | Use |
|---|---|---|
| BicDFw | GCCGAAAAACAGATGGAAGA | *in situ* |
| BicDRv | CGACTCACTATAGGGTCGACATGCTTTGTCTCTGG | *in situ*+ RNAi |
| BicDFwT7 | CGACTCACTATAGGGGCCGAAAAACAGATGGAAGA | RNAi |
