## Supplementary figures and images for "Dynamics of maternal gene expression in *Rhodnius prolixus*"

### Supplementary 8

#### Additional file 8

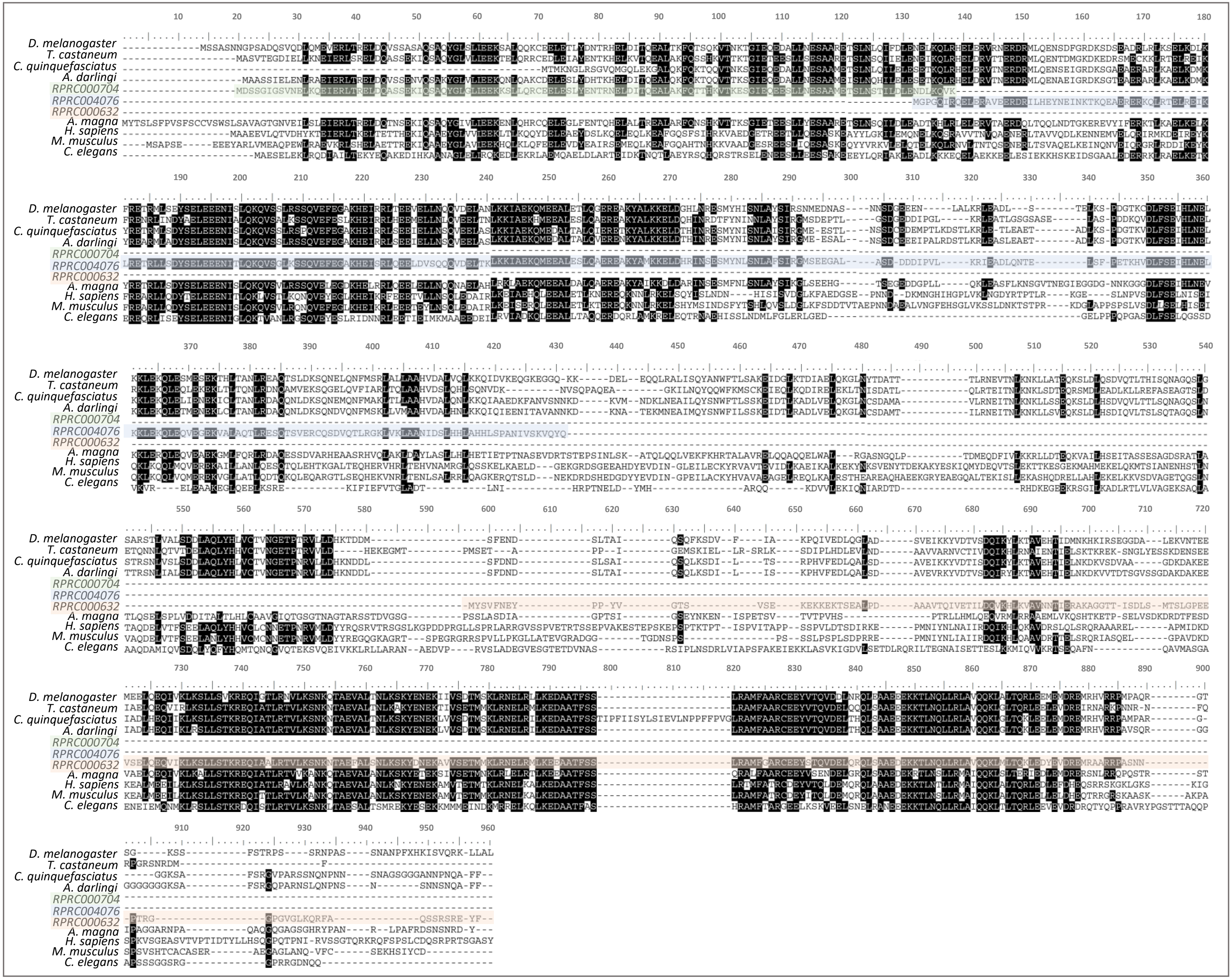
