## Supplementary 9 for "Dynamics of maternal gene expression in *Rhodnius prolixus*"

| Additional file 9: Summary of Parental RNAi experiment. * until the 5th day after injection | | | | | | |
|---|---|---|---|---|---|---|
| **dsRNA** | **dsRNA Dosage** | **Number of injected females** | **% Female survival*** | **Ovipositions** | **Ovary Phenotype** | **Embryo Lethality** |
| **dsRNA*βicD*** | 0,5 μg | 5 | 60% | 22 | 0/3 (0%) | 18/22 (81,8%) |
| | 1,3 μg | 5 | 40% | 9 | 0/2 (0%) | 9/9 (100%) |
| | 2 μg | 6 | 66,60% | 2 | 0/4 (0%) | 2/2(100%) |
| | 2,5 μg | 4 | 50% | 10 | 0/2 (0%) | 10/10(100%) |
| **dsRNA*βlac*** | 1 μg | 2 | 100% | 72 | 0/2 (0%) | 5/72 (6.94%) |
| | 2 μg | 2 | 100% | 148 | 0/2 (0%) | 3/148 (2.02%) |
